# Thermogenic adipocytes alleviate hepatic steatosis and insulin resistance via macrophage cytokine secretion in obese mice

**DOI:** 10.1101/2025.08.01.667482

**Authors:** Hui Wang, Mark Kelly, Najihah Aziz, Emmanouela Tsagkaraki, Sarah M. Nicoloro, Lawrence M. Lifshitz, Adilson Guilherme, Leslie A. Rowland, Batuhan Yenilmez, Kaltinaitis B. Santos, Naideline Raymond, Tiffany DeSouza, Muhammad Sajid, Hanna Choi, Lauren A. Tauer, Jason K. Kim, Silvia Corvera, Michael P. Czech

## Abstract

The *Nrip1* gene encodes the protein Rip140, which suppresses nuclear receptors that regulate energy metabolism. Here we show adipocyte-selective deficiency of Nrip1 (AdNrip1KO) in mice causes a striking expansion of alternatively activated, M2-like macrophages within subcutaneous inguinal adipose tissue (iWAT) in addition to the appearance of thermogenic adipocytes expressing uncoupling protein 1 (UCP1). AdNrip1KO mice are less cold sensitive but showed no differences in whole body energy expenditure or food intake compared to control mice at 22 °C. Strikingly, AdNrip1KO mice on HFD display markedly attenuated hepatic steatosis and insulin resistance compared to control mice on HFD. Secreted factors that might mediate this crosstalk from adipose tissue to liver were searched for by iWAT RNAseq. Unexpectedly, upregulation of genes associated with cytokines and cytokine receptor signaling were the most highly correlated with adipocyte-selective Nrip1 loss in obese mice. Furthermore, the top upregulated genes that encode secreted proteins in AdNrip1KO iWAT are most highly expressed in macrophages, not adipocytes. This list included the IL-1b antagonist IL-1rn, known to attenuate hepatic steatosis and insulin resistance. Indeed, the IL-1rn protein in AdNrip1KO mice was found to circulate at levels we previously reported strongly attenuates hepatic steatosis and glucose tolerance in obese mice. Taken together, these results suggest a paradigm for metabolic crosstalk from thermogenic adipose tissue to liver that is mediated by IL-1rn and potentially other factors secreted from M2-like macrophages within beige adipose tissues.

**Graphic Abstract:** 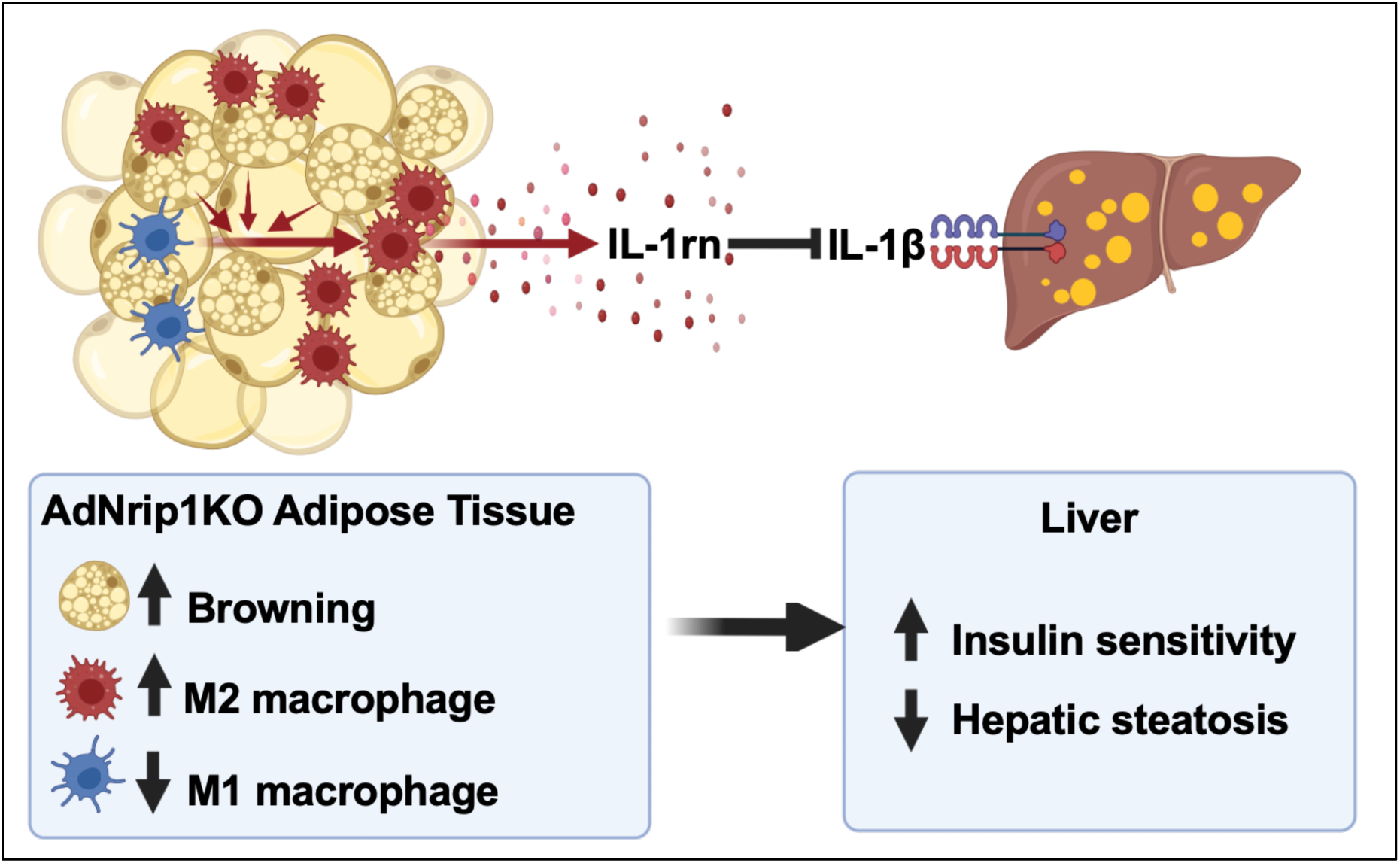

**Highlights:**

- Adipocyte Nrip1 deficiency (AdNrip1KO) strongly promotes adipose browning
- Mouse AdNrip1KO polarizes adipose tissue macrophages towards M2-like
- AdNrip1KO reduces the hepatic fat and insulin resistance of obese mice
- AdNrip1KO adipose tissue macrophages secrete IL-1rn, known to mitigate hepatic fat and insulin resistance

## Introduction

Adipose tissue is a metabolically dynamic organ distributed in many depots throughout the body and plays an essential role in systemic energy and metabolic homeostasis. White adipose tissue (WAT) serves as an energy reservoir, storing excess energy in the form of triglycerides, typically as a single large unilocular lipid droplet per adipocyte (Cinti, 2001; Morigny et al., 2021; Arner & Rydén, 2022). In comparison, brown adipose tissue (BAT) is specialized to dissipate stored chemical energy as heat, protecting mammals against cold stress (Cannon & Nedergaard, 2004; Chouchani et al., 2019). A key player in this cold-induced thermogenesis is mitochondrial uncoupling protein-1 (Ucp1), bypassing ATP production from substrate oxidation (Cannon & Nedergaard, 2011). Morphologically, each single brown adipocyte harbors numerous small lipid droplets (multilocular) and a high density of mitochondria, enabling heat generation (Cannon & Nedergaard, 2011). In addition to the classical brown adipocyte, another population of Ucp1-postive cells, known as beige adipocytes, can be induced within WAT under appropriate pharmacological or physiological stimulations (Timmons et al., 2007; Harms & Seale, 2013; Kajimura et al., 2015). Factors such as cold exposure (Young et al., 1984; Cousin et al., 1993), β-adrenergic agonists (Cousin et al., 1993; Guerra et al., 1998), cardiac natriuretic peptides (Bordicchia et al., 2012), or activation of peroxisome proliferator-activated receptor gamma (PPARγ) (Vernochet et al., 2009; Petrovic et al., 2010) have been shown to powerfully promote beige adipocyte recruitment and/or activate their functions. Other Ucp1-independent thermogenic mechanisms also appear to play important roles in thermogenesis in beige and brown adipocytes, including substrate cycling (Guan et al., 2002; Reshef et al., 2003; Kazak et al., 2015; Brownstein et al., 2022) and calcium cycling (Ikeda et al., 2017). Both WAT and BAT are also endocrine organs, secreting peptides and other mediators that have paracrine and endocrine actions on neighboring cells and distant organs (Harms & Seale, 2013; Villarroya et al., 2017; Ghaben & Scherer, 2019; Czech, 2020).

Increasing evidence has suggested that enhancing the mass and activity of brown or beige fat may hold significant therapeutic potential for countering and reversing obesity, type 2 diabetes (T2D), insulin resistance and other related metabolic pathologies (Poher et al., 2015; Becher et al., 2021). For example, implantation of BAT depots or genetically engineered beige adipocytes has been shown to lower blood glucose and ameliorate fatty liver phenotypes (Stanford et al., 2013; Wang et al., 2020; Tsagkaraki et al., 2021). Specific overexpression of Ucp1 in WAT robustly decreases subcutaneous fat mass and corresponding total body mass compared to their wild-type counterparts (Kopecky et al., 1995). Previous studies indicate Ucp1 abundance is largely dependent on transcriptional regulation, mainly through a promoter region regulated by the cyclic AMP (cAMP) protein kinase A (PKA) signaling pathway and an essential distal enhancer region at the *Ucp1* gene locus (Villarroya et al., 2017; Shapira & Seale, 2019). This enhancer region possesses the response elements that bind key nuclear receptors, including peroxisome proliferator-activated receptors (PPARs) and thyroid hormone receptors (TRs) (Villarroya et al., 2017). Several essential transcriptional regulators have been identified that promote WAT browning, including PPARγ coactivator 1-alpha (Pgc-1α) (Wu et al., 1999) and PR domain-containing protein16 (Prdm16) (Seale et al., 2008). By coactivating PPARγ, Prdm16 transgenic mice promote the conversion of inguinal white adipose cells (iWAT) into beige adipocytes with increased sympathetic innervation in iWAT, rendering their resistance to diet-induced obesity and enhancing energy expenditure (Seale et al., 2011; Cohen et al., 2014).

Conversely, a number of genes have been demonstrated to serve as brakes on *Ucp1* gene expression, including nuclear receptor interacting protein 1 (Nrip1, also known as Rip140) (Leonardsson et al., 2004; Christian et al., 2005; Powelka et al., 2006), Zinc finger protein 423 (Zfp423) (Shao et al., 2016), TLE family member 3 (Tle3) (Villanueva et al., 2013), Twist basic helix-loop-helix transcription factor 1 (Twist1) (Pan et al., 2009) and Interferon regulatory factor 3 (Irf3) (Kumari et al., 2016; Yan et al., 2021). Targeted deletions of these inhibitory transcription factors can limit body weight gain in response to an obesogenic diet by promoting *Ucp1* gene expression in beige adipocytes. We and others previously identified Nrip1 as a transcriptional corepressor of *Ucp1* gene expression, influencing glucose metabolism and lipid oxidation. Nrip1 knockout in adipocytes *in vitro* results in an increase in glucose uptake and lipid oxidation (Christian et al., 2005; Powelka et al., 2006; Kiskinis et al., 2014; Shen et al., 2018; Tsagkaraki et al., 2021). Whole body knockout of Nrip1 in mice display considerable WAT browning with decreased fat mass even on a standard chow diet (Leonardsson et al., 2004). When challenged with a high-fat diet (HFD), Nrip1 null mice display decreased body weight, improved glucose tolerance and reduced liver triglyceride accumulation compared with wild type (WT) controls (Leonardsson et al., 2004). Regardless of diets, deficiency of Nrip1 in mice results in a significantly higher energy expenditure than WT animals, associated with increased oxidative enzymes expression in muscle and intense browning of iWAT and upregulation of Ucp1 (Leonardsson et al., 2004; Seth et al., 2007; Fritah et al., 2012).

Despite the above marked effects of whole body Nrip1 knockout in mice and the thermogenic responses of Nrip1 depletion in adipocytes *in vitro*, tissue specific Nrip1 loss has not been studied for changes in energy metabolism in obese mice. Muscle selective Nrip1KO mice were recently reported, but these mice were not evaluated for glucose tolerance or liver functions (Yamamoto et al., 2023). It is not clear which tissues contribute to the metabolic phenotypes of Nrip1 null mice. Therefore, to determine the contribution of Nrip1/Rip140 expression in adipose tissue to whole body metabolism, we generated an adipose tissue selective Nrip1 knockout mouse (AdNrip1KO). Here, we report these mice reveal a strong axis of regulation between adipose tissue and liver mediated by adipose tissue expression of Nrip1/Rip140. Unexpectedly, AdNrip1KO mice show a striking upregulation of M2-like macrophages in iWAT, accounting for greatly enhanced iWAT expression of macrophage-derived secreted factors. These include the IL-1β antagonist IL-1rn, previously demonstrated by our laboratory and others to attenuate metabolic derangements in livers of obese mice (Petrasek et al., 2012; Negrin et al., 2014).

## Results

### WAT browning in AdNrip1KO mice protects against cold-induced hypothermia

AdNrip1KO mice were generated by crossing Nrip1-floxed mice (Nrip1^flox/flox^) to Adiponectin-Cre animals (Fig. S1A). The qPCR analysis revealed marked disruption of *Nrip1* mRNA expression in interscapular BAT (iBAT), iWAT and gonadal WAT (gWAT), respectively, without detectable effects in liver, skeletal muscle or heart (Fig. 1A). To examine the physiological role of Nrip1, we initially placed mice on a regular chow diet. In contrast to the lean phenotype in whole-body Nrip1-KO mice fed chow diet (Leonardsson et al., 2004), AdNrip1KO mice did not exhibit alterations in body mass, gWAT fat mass (Fig. 1B-C) nor systemic glucose tolerance (Fig. S1B) compared to Nrip1^flox/flox^ controls. However, an enlarged iWAT tissue (Fig. 1D) as well as a trend toward decreased plasma insulin levels (Fig. S1C) was revealed in AdNrip1KO mice.

**Figure 1:**
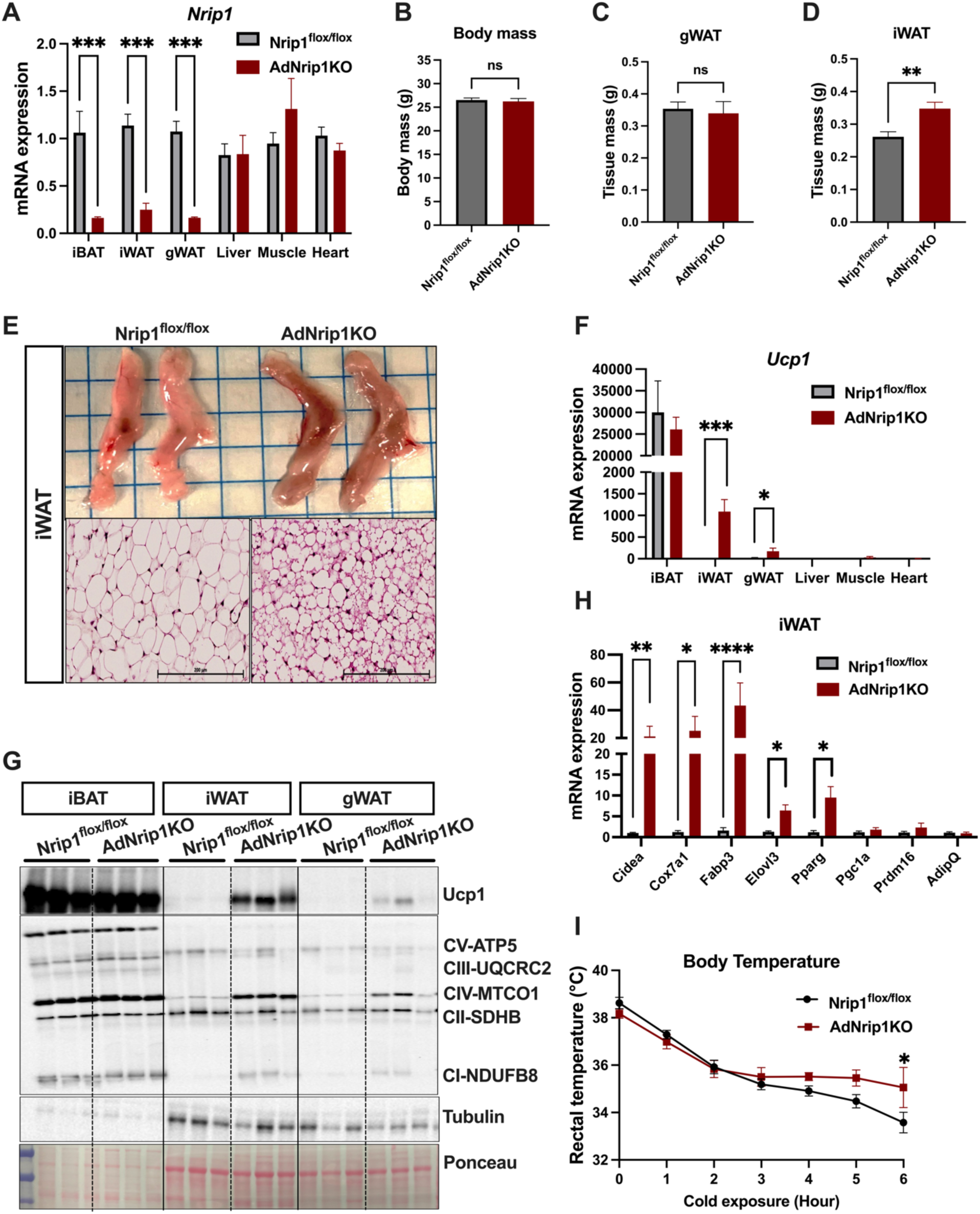
Adipose browning in AdNrip1KO mice with detectable protection from cold-induced hypothermia. **A.** *Nrip1* gene mRNA expression in iBAT, iWAT, gWAT, liver, muscle and heart from 12-week-old male mice fed with chow diet. **B.** Body weight. **C.** gWAT tissue mass. **D.** iWAT tissue mass from the 12-week-old male mice. **E.** Top: macroscopic views of representative iWAT. Below: microscopic images of representative iWAT stained with Hematoxylin and Eosin. **F.** Quantitative PCR analysis of *Ucp1* gene expression in iBAT, iWAT, gWAT, liver, muscle and heart from 12-week-old male mice fed with chow diet. **G**. Western blot analysis of Ucp1 protein abundance and mitochondrial oxidative phosphorylation proteins in iBAT, iWAT and gWAT. **H.** Quantitative PCR analysis of thermogenic genes expression in iWAT. **I.** Rectal body temperature during acute cold exposure without food on 12-week-old male mice. All male mice in these studies were fed with chow diet, n = 12 mice per genotype. Data are represented as mean ± SEM. In panels **A**, **F**, **H** and **I**, comparisons were performed by Two-way ANOVA with Sidak’s multiple comparisons test. In data **B**, **C**, **D**, statistical analyses were compared with unpaired 2-tailed *t*-test between Nrip1^flox/flox^ vs AdNrip1KO animals. ns, not significant, \**p* < 0.05, \*\**p* < 0.005, \*\*\**p* < 0.0005, \*\*\*\**p* < 0.0001.

Notably, AdNrip1KO iWAT showed a deeper brown color than their Nrip1^flox/flox^ controls (Fig. 1E). Hematoxylin and eosin (H&E) staining revealed numerous smaller lipid droplets in a multilocular pattern in the adipocytes of AdNrip1KO iWAT, consistent with the profound browning effects (Fig. 1E). Multilocular adipocytes were also found in gWAT of AdNrip1KO, but to a lesser degree than that in iWAT (Fig. S1D). As expected, Ucp1 mRNA and protein expression were upregulated in both iWAT and gWAT of AdNrip1KO animals (Fig. 1F-G). Mitochondrial oxidative phosphorylation (OXPHOS) proteins in white fat depots of AdNrip1KO were increased dramatically as well, suggesting enhanced mitochondrial biogenesis (Fig. 1G). Furthermore, expression of many other thermogenic marker genes in iWAT of AdNrip1KO mice were also significantly increased, including the higher expression of *Cidea* (Cell death inducing DNA fragmentation factor alpha-like effector A), *Cox7a1* (Cytochrome C Oxidase Subunit 7A1), *Fabp3* (Fatty acid binding protein 3), *Elovl3* (ELOVL fatty acid elongase 3) and *PPARγ* (Fig. 1H). However, no notable alterations in iBAT between groups were observed related to Ucp1 expression, mitochondrial protein contents, nor morphological phenotypes (Fig. 1G and S1D). To further assess the functional implications of Ucp1 induction, mice were exposed to an acute cold acclimation for 6 hours without food access. Although both Nrip1^flox/flox^ and AdNrip1KO mice showed gradually decreasing rectal temperatures under cold exposure, AdNrip1KO mice had the ability to maintain a relatively higher body temperature at 6 hours than their Nrip1^flox/flox^ littermates (Fig. 1I).

Primary adipocyte cell cultures derived from preadipocytes in adipose tissues isolated from iWAT of AdNrip1KO mice displayed higher Ucp1 expression than the cultures from Nrip1^flox/flox^ littermates when fully differentiated (Fig. S1E-F), confirming the cell-autonomous effects of Nrip1 on *Ucp1* expression. Taken together, these findings suggest that Nrip1 deficiency in adipocytes profoundly promotes Ucp1 expression both *in vivo* and *in vitro*, associated with protection from cold-induced hypothermia.

### AdNrip1KO mice display improved insulin sensitivity independent of body mass

To further examine the metabolic changes in AdNrip1KO mice, they were challenged with a HFD (60% Cal from fat). Compared to Nrip1^flox/flox^ controls, HFD fed AdNrip1KO mice exhibited a significant improvement in glucose tolerance (Fig. 2A). Whole-body insulin sensitivity was also increased in AdNrip1KO animals, as determined by insulin tolerance tests (Fig. 2B). Although the body weight was comparable between groups (Fig. 2C), iBAT of AdNrip1KO mice showed a marked reduction in tissue mass (Fig. 2D). Histological analysis revealed that prolonged HFD feeding resulted in massive fat accumulation in iBAT of Nrip1^flox/flox^ mice, compared with the age-matched Nrip1^flox/flox^ mice maintained on chow diet. However, this effect was partially alleviated in AdNrip1KO mice on a HFD (Fig. 2D).

**Figure 2:**
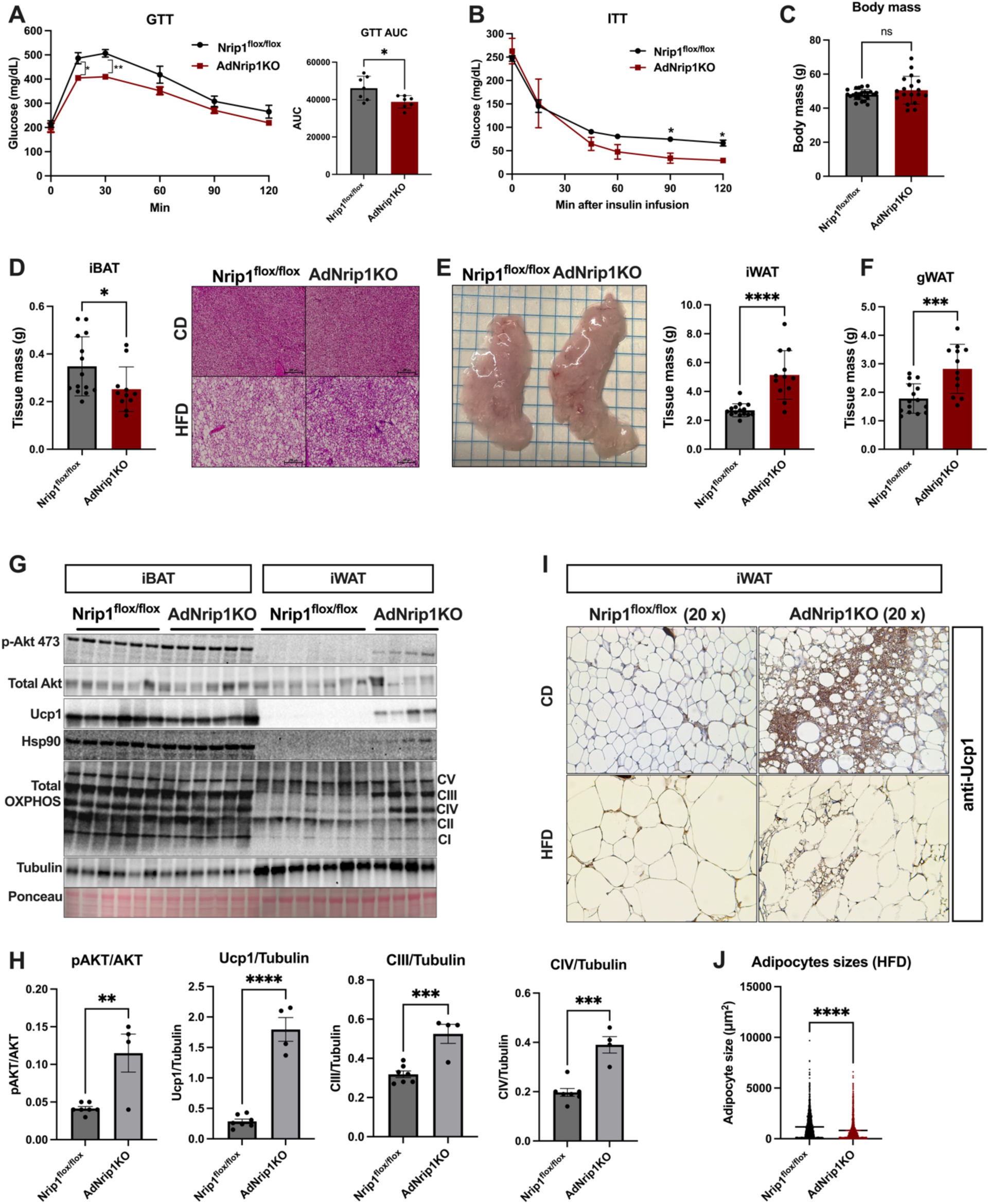
AdNrip1KO mice display improved insulin sensitivity independent of body mass. **A.** Left: Glucose tolerance test (GTT) during HFD feeding. Right: area under the curve quantification of GTT results. **B.** Insulin tolerance test (ITT) during HFD feeding. **C.** Body mass after 16 weeks HFD feeding. **D.** iBAT tissue mass and the H&E staining of iBAT tissue from age-matched mice fed either chow or HFD. **E.** Left, macroscopic views of representative iWAT after HFD feeding. Right: iWAT tissue mass after 16 weeks HFD feeding. **F.** gWAT tissue mass. **G.** Protein abundance analysis of Ucp1, p-Akt473, total Akt, Hsp90 and mitochondrial oxidative phosphorylation proteins in iBAT and iWAT from mice after 16 weeks HFD feeding. **H.** Protein abundance quantification from G. **I.** Ucp1 antibody immunohistochemical staining of iWAT. n = 11-14 mice per genotype for animal feeding experiment. **J.** Quantification of adipocyte size, 2024 and 1379 adipocytes were quantified from Nrip1^flox/flox^ and AdNrip1 iWAT, respectively. Data are represented as mean ± SEM. **A left** and **B** comparisons were performed by Two-way ANOVA with Sidak’s multiple comparisons test. In data **A right**, **C**, **D left**, **E right**, **F, H and J** statistical analysis were compared with unpaired 2-tailed *t*-test between Nrip1^flox/flox^ vs AdNrip1KO animals. ns, not significant, \**p* < 0.05, \*\**p* < 0.005, \*\*\**p* < 0.0005, \*\*\*\**p* < 0.0001.

An unexpected result in these studies was the increased mass of iWAT and gWAT depots in AdNrip1KO mice, particularly in iWAT (Fig. 2E-F). Despite this adipose expansion, *Ucp1* mRNA expression remained higher in iWAT and gWAT of AdNrip1KO mice than corresponding tissues in Nrip1^flox/flox^ mice (Fig. S2A-F). Mitochondrial OXPHOS proteins and Ucp1 protein abundance were elevated in iWAT of AdNrip1KO mice (Fig. 2G). Also, dramatically higher amounts of phosphorylated Akt/protein kinase B at serine 473 (pAkt-S473) in iWAT were observed in AdNrip1KO, suggesting their enhanced insulin signaling sensitivity (Fig. 2G-H). The molecular chaperone Hsp90, which regulates PPARγ stability and adipocyte differentiation, was increased in AdNrip1KO mice (Fig. 2G-H). Consistently, immunohistochemical analysis revealed abundant Ucp1-positive signals in the iWAT of AdNrip1KO mice maintained on a chow diet (Fig. 2I). After prolonged HFD feeding, Ucp1-expressing multilocular fat cells were still present in the iWAT of AdNrip1KO mice, but not in Nrip1^flox/flox^ mice (Fig. 2I). Comparison of adipocyte sizes in iWAT between Nrip1^flox/flox^ and AdNrip1KO mice revealed the smaller adipocytes in AdNrip1KO animals (Fig. 2J). Given considerable enlargement of the iWAT mass and size in AdNrip1KO mice, the smaller beige adipocytes suggest that the increase in fat mass is due to adipocyte hyperplasia rather than hypertrophy.

### AdNrip1KO mice show marked improvements in hepatic insulin responsiveness

To investigate the systemic insulin sensitivity, hyperinsulinemic-euglycemic clamp studies were conducted in AdNrip1KO mice after 16 weeks of HFD. Basal glucose levels were similar between Nrip1^flox/flox^ mice and AdNrip1 KO mice (Fig. 3A). The plasma glucose levels were maintained at 109 ± 7 mg/dL and 116 ± 9 mg/dL in Nrip1^flox/flox^ and AdNrip1KO animals, respectively, during the 2-hour clamps. The steady-state glucose infusion rates required to maintain euglycemia were profoundly increased by 3-fold to 24.1 ± 1.4 mg/kg/min in AdNrip1KO mice compared to 8 ± 1.8 mg/kg/min in Nrip1^flox/flox^ mice, indicating greatly improved insulin sensitivity in HFD-fed AdNrip1KO mice (Fig. 3B). Whole-body glucose turnover and glycogen synthesis were also increased in AdNrip1KO mice compared to Nrip1^flox/flox^ littermates (Fig. 3C-D), suggesting improved peripheral glucose metabolism in AdNrip1KO mice. Insulin-stimulated glucose uptake in brown adipose tissue and skeletal muscle of AdNrip1KO mice trended higher in AdNrip1KO mice compared to Nrip1^flox/flox^ mice (Fig. 3E-F).

**Figure 3:**
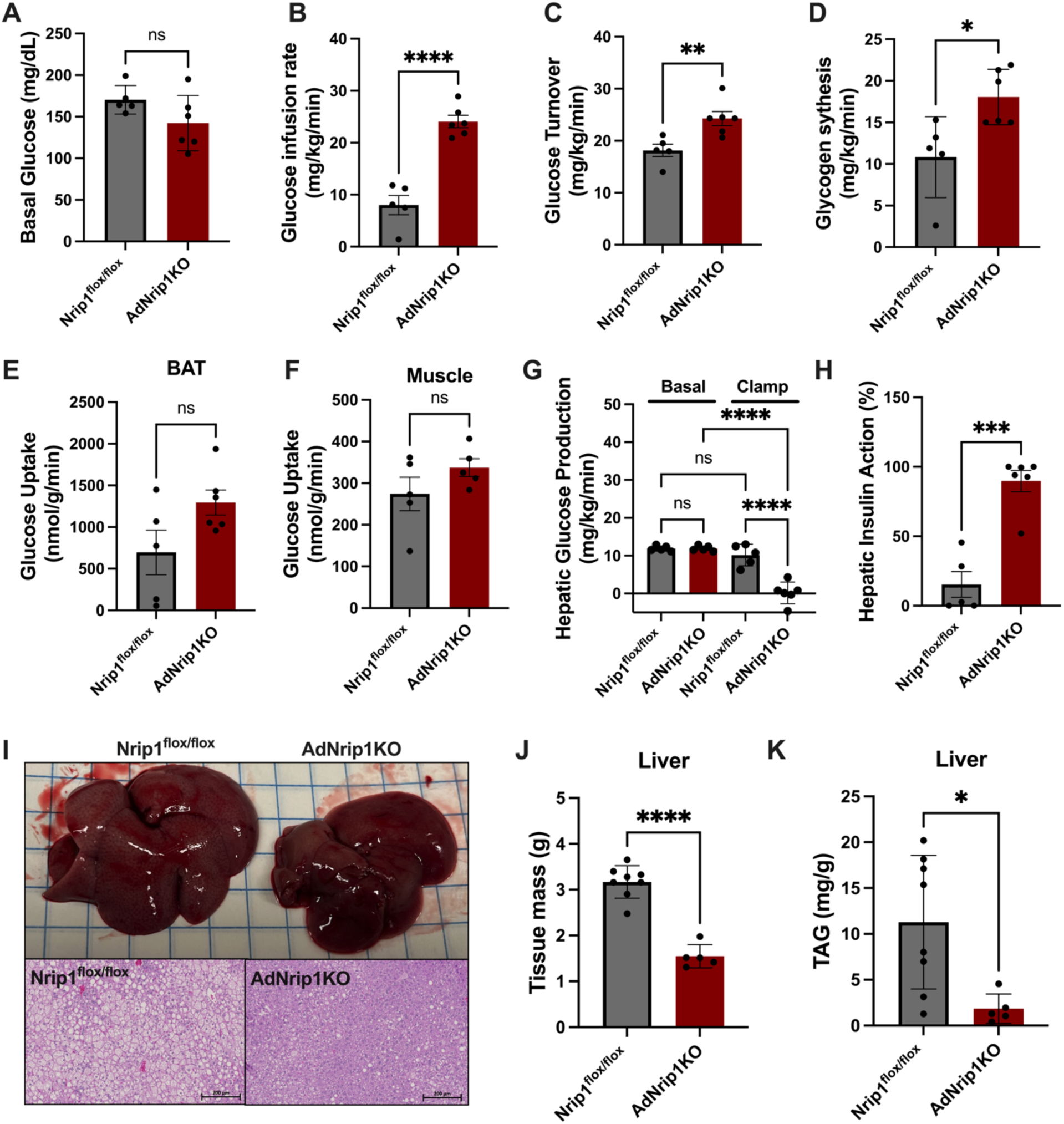
AdNrip1KO improves hepatic metabolism during hyperinsulinemic-euglycemic clamp conditions. **A.** Basal glucose levels after 16 weeks HFD feeding. **B**. Glucose infusion rate to reach the steady state phase of the hyperinsulinemic-euglycemic clamp study after 16 weeks HFD feeding. **C.** Whole-body glucose turnover rates. **D.** Whole-body glycogen synthesis. **E-F.** Glucose utilization in iBAT and muscle. **G.** Hepatic glucose production in basal and clamp conditions for both genotypes. **H.** Hepatic insulin action. **I.** Top, macroscopic views of representative liver from Nrip1^flox/flox^ and AdNrip1 mice. Below, microscopic images of representative liver with Hematoxylin and Eosin staining. **J.** Liver tissue mass and **K.** Liver Triglyceride content. All mice were fed with HFD for 16 weeks before clamp study (n = 6 for Nrip1^flox/flox^ mice, n = 5 for AdNrip1KO mice). Data are represented as mean ± SEM. In panel **G**, comparisons were analyzed by One-way ANOVA with Tukey’s multiple comparisons test. All other bar chart statistics were analyzed with unpaired 2-tailed *t*-test between Nrip1^flox/flox^ vs AdNrip1KO animals. ns, not significant, \**p* < 0.05, \*\*\**p* < 0.0005, \*\*\*\**p* < 0.0001.

Basal hepatic glucose production (HGP) was similar between groups, but insulin infusion caused a minimal suppression of HGP during clamps in Nrip1^flox/flox^ mice, indicating severe insulin resistance in the liver after HFD (Fig. 3G). Strikingly, insulin-mediated suppression of HGP was increased and almost completely restored in AdNrip1KO mice, demonstrating improved hepatic insulin action in HFD-fed AdNrip1KO mice (Fig. 3G-H). Additionally, we observed that liver weights of AdNrip1KO mice were approximately 50% of Nrip1^flox/flox^ mouse livers (Fig. 3I-J). Histological investigation found significantly less lipid accumulation in liver of AdNrip1KO mice, with a marked reduction in liver triglyceride (TAG) content (Fig. 3K).

To further investigate glucose uptake in adipose tissues, a separate cohort of 16wks-HFD fed mice were analyzed using an *ex vivo* glucose uptake technique. For this assessment, iBAT, iWAT, and gWAT were collected and incubated in glucose-free DMEM, followed with exogenous application of 5 mM glucose at 37**°**C. Media was collected 0, 2, 4, 6, and 8 hours and monitored for glucose to assess glucose utilization in these *ex vivo* tissues. Notably, while glucose uptake in iBAT was comparable between the two genotypes, higher glucose uptake was observed in iWAT and gWAT of AdNrip1KO mice (Fig. S2G-I).

To better understand the underlying mechanisms accounting for the above improved liver insulin sensitivity, we hypothesized that the adipose thermogenesis in AdNrip1KO mice may reduce circulating free fatty acids (FFAs) to protect mice from liver steatosis (Perry et al., 2015; Shum et al., 2016; Titchenell et al., 2016). To test this, we quantified circulating FFA levels under fasting and glucose-stimulated conditions and found there were no significant differences in plasma insulin, FFA or TAG levels between AdNrip1KO and Nrip1^flox/flox^ animals (Table 1). However, significant decreases in plasma total cholesterol, high-density lipoprotein (HDL), and low-density lipoprotein (LDL) in fasted AdNrip1KO mice were noted.

**Table 1.** Plasma factors in Nrip1^flox/flox^ and AdNrip1KO mice after HFD feeding.

|  | Nrip1flox/flox | AdNrip1KO |
| --- | --- | --- |
| Fasting Insulin (μIU/mL) | 47.25 ± 7.48 | 39.77 ± 4.02 |
| Refed Insulin (μIU/mL) | 74.14 ± 17.99 | 75.79 ± 1.65 |
| Fasting Plasma FFA (μM) | 240.2 ± 237.8 | 478.1 ± 142.9 |
| Refed Plasma FFA (μM) | 533.1 ± 61.50 | 618.4 ± 85.31 |
| Fasting total Triglyceride (mg/dL) | 88.33 ± 9.58 | 89.00 ± 0.67 |
| Fasting total Cholesterol (mg/dL) | 206.30 ± 17.83 | 151.70 ± 9.02 **** |
| Fasting HDL (mg/dL) | 156.30 ± 16.30 | 124.80 ± 14.80*** |
| Calculated Fasting LDL (mg/dL) | 35.67 ± 10.45 | 11.00 ± 5.00** |
| Calculated Fasting VLDL (mg/dL) | 17.67 ± 5.27 | 17.80 ± 1.80 |
| Fasting Gdf15 (pg/mL) | 90.90 ± 21.91 | 117.78 ± 16.33 |
| Fasting Adiponectin (μg/mL) | 3.797 ± 0.3547 | 2.215 ± 0.3224**** |

After 14 weeks HFD feeding, mice were fasted overnight with free access to water. The next morning, fasting blood samples were collected for quantification. After D (+) glucose administration (intraperitoneal injection, 1g/kg body weight), blood samples were collected again for quantification. Results are expressed as means ± SE (n = 5-8), compared with unpaired 2-tailed *t*-test between Nrip1<sup>flox/flox</sup> vs AdNrip1KO animals, \**p* < 0.05, \*\**p* < 0.005, \*\*\**p* < 0.0005, \*\*\*\**p* < 0.0001.

### AdNrip1KO mice display normal energy expenditure when housed at 22 °C

Given the increased energy expenditure in whole-body Nrip1 null mice and the higher Ucp1 protein abundance in adipose tissues in AdNrip1KO mice, indirect calorimetry using metabolic cages was performed in mice after 14 weeks of HFD. However, there were no distinguishable differences between AdNrip1KO mice and controls in oxygen consumption rates (Fig. 4A), respiratory exchange ratio (Fig. 4B), or energy expenditure (Fig. 4C-D). Food and water intake also showed no significant differences between the two groups of mice (Fig. 4E-F). Notably, AdNrip1KO mice displayed an overall higher locomotor activity than their Nrip1^flox/flox^, especially during light times (Fig. 4G-H).

**Figure 4:**
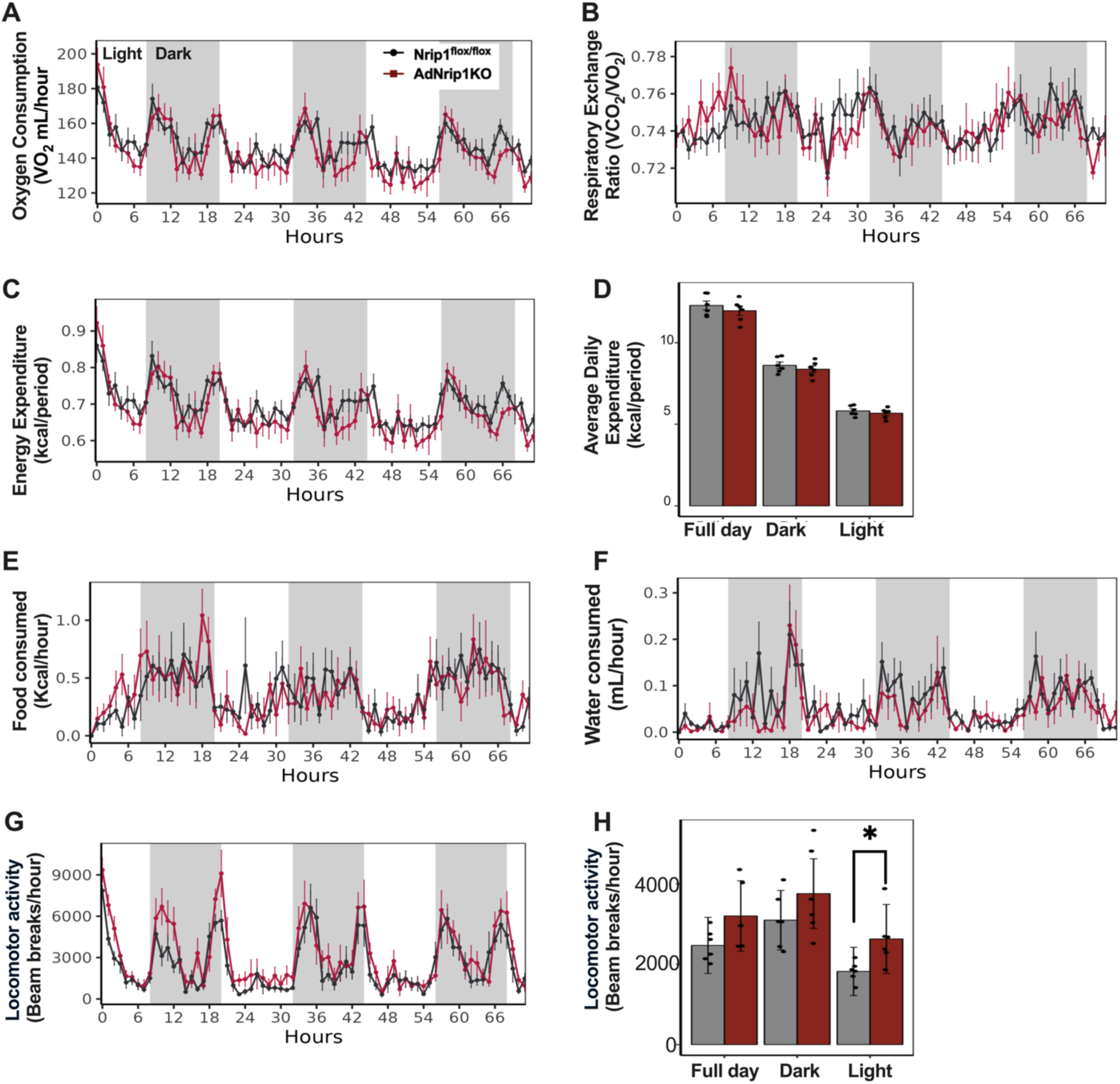
AdNrip1KO mice do not display detectable increases in energy expenditure during HFD feeding. **A.** Average oxygen consumption by indirect colorimetry for a consecutive 64-hour period monitoring in metabolic cages. **B.** Respiratory exchange ratio. **C.** Energy expenditure. **D.** Average daily energy expenditure quantification. **E.** Food consumption and **F**. Water intake**. G.** Locomotor activity, and **H**. the corresponding average daily locomotor activity quantification (n = 6 mice for each genotype). The metabolic quantification experiment was performed after 14 weeks HFD feeding at room temperature. Data are represented as mean ± SEM. All data are analyzed using CalR software, while the total body mass used as selective covariant. Statistical analyses were compared by Two-way ANOVA with Sidak’s multiple comparisons test. \**p* < 0.05.

### AdNrip1KO iWAT contains an increased abundance of M2-like macrophages

To characterize the AdNrip1KO mice in an unbiased manner, transcriptome analysis was performed by RNA-sequencing in iWAT from Nrip1^flox/flox^ and AdNrip1KO mice after 16 weeks of HFD feeding. Principle component analysis from the two groups revealed a clear separation into distinct clusters within samples, suggesting a distinct gene expression pattern between the two genotypes (Fig. 5A). Differential gene expression analysis revealed 1222 significantly upregulated genes (cutoff padj<0.05, FC ≥ 2) in iWAT of AdNrip1KO mice, including a 72-fold increase in *Akrlc1 (Aldo-keto reductase family 1 member C-like)*, 65.8-fold increase in *Ucp1*, and 27.5-fold increases in *Nptx1 (Neuronal pentraxin 1)* expression (Fig. 5B and Table S1). Meanwhile, 1377 genes with significantly decreased expression were identified including *Nrip1*, *Retn (Resistin)* and *Rbp4 (Retinol-binding protein 4)* (Fig. 5B). As expected, many genes encoding mitochondrial proteins were substantially upregulated in the AdNrip1KO iWAT. These include genes involved in the mitochondrial respiratory chain (*Cox7a1, Cytochrome C oxidase subunit 7A1; Atp8b4, ATPase H+ Transporting Multi-subunit Proton Pump; Cox8b, Cytochrome C oxidase subunit 8B*), fatty acid oxidation (*Cptb1b, Carnitine palmitoyltransferase 1B; ChkbCptb1b, Choline kinase-like Carnitine palmitoyltransferase 1B*), and mitochondrial biogenesis (*Mtfr2, Mitochondrial fission regulator 2; Vat1, Vesicle amine transport 1*) (Fig. 5C). Additionally, genes controlling lipid synthesis (*Srebf1, Sterol regulatory element binding transcription factor 1; Irs2, Insulin receptor substrate 2; Fasn, Fatty acid synthase; Dgat 2, Diacylglycerol O-acyltransferase 2*) and lipid droplet formation (*Cidec, Cell death inducing DFFA Like effector C*), were decreased dramatically while genes regulating lipid oxidation were strongly enhanced (*PPARa*, *Peroxisome proliferator-activated receptor Alpha*; *Pdk4*, *Pyruvate dehydrogenase kinase 4*; *Aldh1l2, Aldehyde dehydrogenase 1 family member L2*) (Fig. 5D).

**Figure 5:**
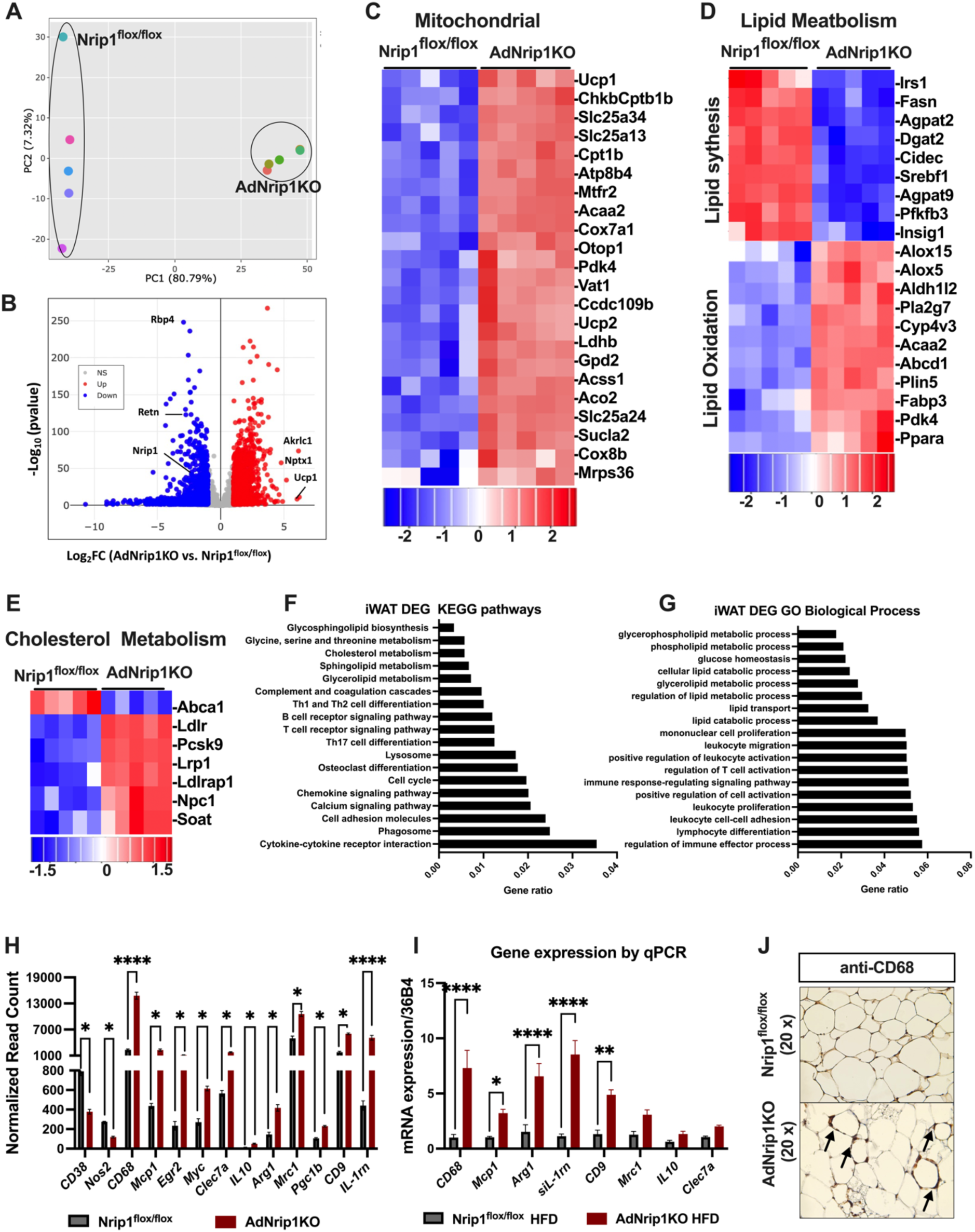
RNA sequencing analysis on iWAT of Nrip1^flox/flox^ and AdNrip1KO mice after HFD feeding. **A.** Principal component plot prior to differential gene expression (DEG) analysis, (n = 5 mice/genotype). **B.** Volcano plot of DEGs, upregulated genes in AdNrip1KO were marked with Red dot, downregulated in Blue dot, with data analysis cutoff FC≥2, padj<0.05. **C**. Upregulated mitochondrial genes of AdNrip1KO iWAT. **D.** DEGs regulating lipid metabolism between Nrip1^flox/flox^ and AdNrip1KO groups. **E.** DEGs regulating cholesterol metabolism. **F.** KEGG pathway analysis of differentially expressed genes, padj<0.05. **G.** Enriched GO term-biological process based on the differentially expressed genes, padj<0.05. **H.** Normalized read counts of macrophage genes by RNA sequencing, \**p* < 0.05, \*\*\*\**p* < 0.0001. **I.** Quantitative PCR analysis of gene expression in iWAT of Nrip1^flox/flox^ and AdNrip1KO mice after 16-weeks HFD feeding. Data are represented as mean ± SEM. **J.** Cd68 antibody immunohistochemical staining of iWAT. **H** and **I** Statistical analyses were compared by Two-way ANOVA, \**p* < 0.05, \*\**p* < 0.005, \*\*\*\**p* < 0.0001 (n = 5 -7 mice/genotype).

To explore the basis for the decreased levels of circulating total cholesterol in AdNrip1KO mice, we compared the expression of key genes related to cholesterol metabolism. Interestingly, we found that genes regulating cholesterol uptake were significantly upregulated in iWAT of AdNrip1KO mice, particularly *Ldlr* (*Low-density lipoprotein receptor*) and *Ldlrap1* (*Low-density lipoprotein receptor adaptor protein 1*). In contrast, the cholesterol efflux gene *Abca1* (*ATP-binding cassette transporter A1*), which plays a critical role in transporting cholesterol from cells to the bloodstream, was dramatically decreased in AdNrip1KO iWAT (Fig. 5E). These data suggest enhanced cholesterol uptake and utilization in AdNrip1KO iWAT may in part explain the changes in circulating cholesterol.

Next, enriched KEGG pathway and GO-Term biological process analyses on differentially expressed genes in Nrip1^flox/flox^ versus AdNrip1KO iWAT were performed. Unexpectedly, a large number of regulated genes by Nrip1KO in adipocytes were found to be related to chemokine-chemokine receptor signaling, immune cell receptor signaling pathways, lymphocyte differentiation and immune effector process followed by alterations in lipid and cholesterol metabolism (Fig. 5F-G). Notably, the normalized read counts from the RNA sequencing showed dramatically altered gene expression related to adipose tissue macrophage (ATMs) recruitment (*Mcp1 (Monocyte chemoattractant protein-1)* and *CD68 (CD68 Molecule)*), with decreased classical activated M1-like macrophage markers (*CD38 (CD38 Molecule)* and *Nos2 (Nitric oxide synthase 2))*. In contrast, robust increases in expression of alternatively activated M2-like macrophage markers such as *Arg1 (Arginase 1)*, *IL-1rn* (*Interleukin-1 receptor antagonist*),*CD9 (CD9 Molecule)*, *Mrc1 (Mannose receptor C-Type 1, or CD206)*, *Egr2 (Early growth response 2)*, *Myc (MYC proto-oncogene, BHLH Transcription Factor)*, *Clec7a (C-Type Lectin Domain Family 7 Member A)*, *IL10 (Interleukin 10)*, *Ppargc1b (PPAR*γ *coactivator 1 Beta)* were observed, and these changes were validated by mRNA expression qPCR quantification (Fig. 5H-I) (Gordon, 2003; Martinez et al., 2006; Stienstra et al., 2008). Consistent with these findings, immunohistochemical analysis revealed the crown-like structures with anti-CD68 staining around adipocytes confirmed the significantly increased CD68 presence in AdNrip1KO iWAT samples (Fig. 5J). Taken together, these data indicate that adipocyte Rip140 not only controls Ucp1 expression and thermogenesis in adipocytes *in vivo*, but also modulates the cellular composition and microenvironment in the beige adipose tissue, including a marked expansion of M2-like macrophages.

### Enhanced circulating IL-1rn in AdNrip1KO mice

The large increase in abundance of CD68-positive, M2-like macrophages in iWAT of AdNrip1KO mice raised the hypothesis that the dramatically decreased liver triglyceride and increased insulin sensitivity in AdNrip1KO mice may be mediated by factors secreted by adipose tissue macrophages rather than the Nrip1KO adipocytes themselves. To test this, ∼130 upregulated genes annotated for encoding secreted proteins were identified in the RNAseq data form iWAT in control and AdNrip1KO mice (Table S1) using the online DAVID bioinformatics tool (https://davidbioinformatics.nih.gov/) (Sherman et al., 2022) and UniProt (https://www.uniprot.org). In order to restrict the large number in this list, we took advantage of a similar RNAseq database derived from our published experiments (Tsagkaraki et al., 2021, 2023) in which Nrip1KO mouse adipocytes are implanted into mice. Such implanted cells grow into adipose tissue depots that also cause beneficial liver phenotypes (Tsagkaraki et al., 2021). As shown in Figure 6A, comparison between the RNAseq data from Figure 5 with the previously published data on the implants (Tsagkaraki et al., 2021, 2023) identified 61 common upregulated genes predicted to encode secreted proteins (Table S2). Applying more stringent criteria (adjusted p-value < 0.05, fold change ≥ 5, normalized RNA sequencing counts ≥ 200), we narrowed the list to 20 candidate genes of highest interest (Fig. 6A and Table S2). Consistent with the hypothesis, all of these genes were much more highly expressed in macrophages than adipocytes, according to the online Single Cell Portal ((https://singlecell.broadinstitute.org/single_cell) (Emont et al., 2022) (Fig. S3 and Table S2). Among them, *IL-1rn* and *Gdf15* (*Growth differentiation factor 15*) exhibited more than 10-fold increases in expression in AdNrip1KO iWAT according to these data sets, with particularly abundant expression levels for *IL-1rn* (Tables S1, S2).

**Figure 6:**
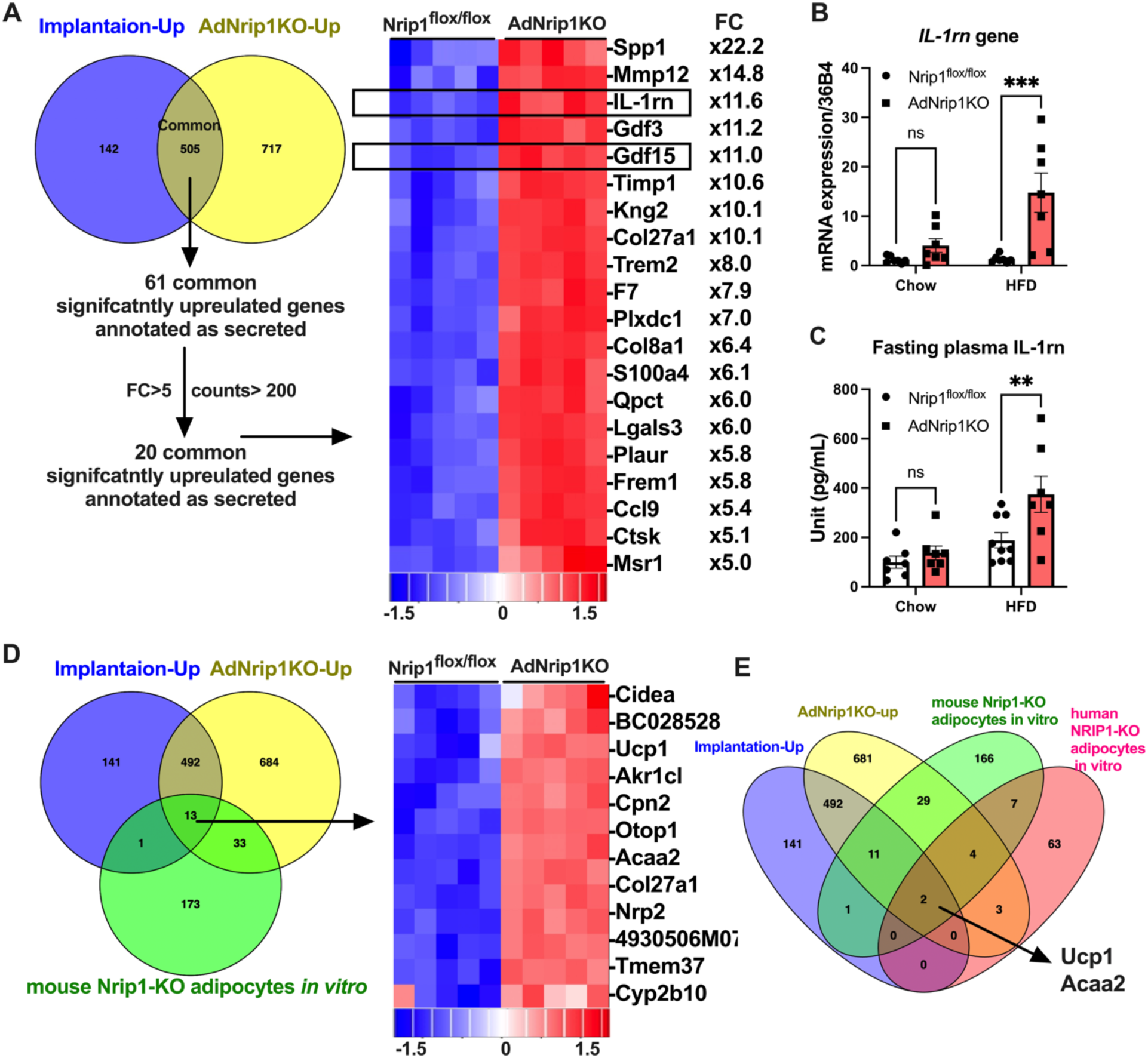
Increased circulating IL-1rn protein associated with expansion of M2-ike ATMs in AdNrip1KO mice. **A.** Venn diagram revealed the common upregulated gene expression between Nrip1KO implanted samples and AdNrip1KO iWAT, which further identified common secreted proteins. **B.** mRNA expression validation of secreted IL-1rn by qPCR. **C.** ELISA quantification of plasma secreted IL-1rn in circulation. Data are represented as mean ± SEM, Statistical analyses were compared by Two-way ANOVA with Sidak’s multiple comparisons test (n = 7 - 9 /group). **D.** Venn diagram analysis of upregulated DEGs among two *in vivo* Nrip1KO mouse models and the *in vitro* mouse Nrip1KO adipocytes. **E.** Venn diagram analysis of upregulated DEGs among the *in vivo* Nrip1ko mouse models, *in vitro* mouse Nrip1KO adipocytes and in vitro human NRIP1KO adipocytes.

Our group and others have previously shown that *in vivo* administration of IL-1rn (Negrin et al, 2014) or Gdf15 (Sjøberg et al., 2023) reduces hepatic lipid accumulation and improves systemic glucose tolerance. Consistently, qPCR quantification confirmed that mRNA expression of the secreted isoform of *IL-1rn* (*sIL-1rn*) was strikingly increased in AdNrip1KO iWAT from HFD fed mice (Fig. 6B). Therefore, we next tested whether adipose selective loss of Nrip1/Rip140 leads to increased levels of circulating IL-1rn and Gdf15 in lean and obese mice. Blood levels of IL-1rn were not affected in AdNrip1KO mice compared to Nrip1^flox/flox^ control mice on a chow diet, but a significant two-fold increase in circulating IL-1rn peptide was observed in obese AdNrip1KO mice fed HFD (Fig. 6C) compared to control mice on HFD. In contrast, circulating Gdf15 levels showed no significant difference in AdNrip1KO versus control mice on HFD (Table 1). These data are consistent with the concept that increased M2-like macrophages in the beige adipose tissue of AdNip1KO mice secrete elevated levels of IL-1rn, which in turn regulates hepatic lipid accumulation and insulin-sensitive glucose production.

Finally, we considered whether any of the mRNAs of the candidate secreted proteins in adipose tissue from AdNrip1KO mice might be upregulated in primary mouse adipocytes when depleted of Nrip1 *in vitro*. However, only 13 genes were identified as commonly upregulated genes across all three datasets, and only the matrix protein Col27a1 of the secreted proteins identified in AdNrip1KO adipose tissue were among them (Fig. 6D). Seven of these genes are recognized as classical cold-induced browning markers (Rosell et al., 2014), such as *Ucp1*, *Cidea*, *Acaa2* (*Acetyl-CoA acyltransferase 2*)*, Otop1* (*Otopetrin 1*), *Cpn2* (*Carboxypeptidase N subunit 2*) *and Tmem37* (*Transmembrane Protein 37*). Although the enzyme Cpn2 can also be secreted into circulation, it showed minimal expression in adipose tissues (Table S1). When we extended this type of analysis to include RNAseq data from human NRIP1KO adipocytes *in vitro,* only two genes—*Ucp1* and *Acaa2*—consistently showed upregulation across all four datasets (Fig. 6E).

## Discussion

Previous findings showed that the single nucleotide polymorphisms in the human *Nrip1* gene locus are associated with multiple metabolic phenotypes (https://hugeamp.org/gene.html?gene=NRIP1) (Dornbos et al., 2022) and that HFD fed Nrip1 null mice remain lean with a metabolically healthy physiology (Leonardsson et al., 2004; Powelka et al., 2006; Seth et al., 2007). These findings raised a key question in the field: which tissues that express Nrip1/RIP140 contribute to the remarkable systemic effects elicited by its genetic deletion? To date, tissue selective Nrip1KO mice have not previously been characterized in sufficient detail to address this question. Liver-selective knockdown of Nrip1/Rip140 using adenovirus-mediated expression of shRNA revealed increased lipoprotein release and decreased steatosis, but HFD feeding, glucose tolerance and energy expenditure were not studied under these conditions (Diaz et al., 2008). Likewise, genetic deletion of Nrip1/Rip140 in striated muscle and cardiomyocytes revealed important effects on cardiovascular function in mice (Yamamoto et al., 2023), but detailed metabolic analysis was not reported. Since increased expression of genes related to thermogenesis, including the uncoupler protein Ucp1, is a common feature of both adipose tissue in the Nrip1 null mouse (Leonardsson et al., 2004) and in cultured adipocytes *in vitro* (Christian et al., 2005; Powelka et al., 2006; Kiskinis et al., 2014; Shen et al., 2018; Tsagkaraki et al., 2021), we asked whether AdNrip1KO mice might phenocopy Nrip1 null mice. A major overall conclusion of the data presented here is that these two mouse models do share the beneficial metabolic features of increased glucose tolerance, enhanced insulin sensitivity and reduction of liver triglyceride. However, the AdNrip1KO mice do not exhibit the high energy expenditure and maintenance of lean body weight on HFD as observed in the whole body Nrip1KO mice.

The experiments reported here demonstrate a strong axis of metabolic interorgan communication between adipose tissue and liver. The strongly decreased hepatic steatosis (Fig 3I-K) and improved hepatic insulin action (Fig. 3G-H) in the HFD-fed AdNrip1KO mice compared to control mice are striking. This is especially impressive in the context of no change in overall body weight (Fig. 2C) and increases in the mass of inguinal and gonadal adipose depots in these mice (Fig. 2E-F). The conventional view of adipose tissue-directed effects on the liver is that fatty acids and glycerol released from adipocytes are main factors that modulate hepatic glucose production (Samuel & Shulman, 2012; Czech, 2017). Thus, lipolytic activity in adipose tissue releases fatty acids that are esterified to triglyceride in hepatocytes, contributing to steatosis and lipoprotein synthesis. Glycerol, the other product of lipolysis, serves as a substrate for liver gluconeogenesis. Knockout or pharmacological inhibition of adipose triglyceride lipase (Atgl), which reduces FFAs released from WAT, decreases WAT-derived hepatic acetyl-CoA levels, thereby protecting mice from high-fat diet-induced insulin resistance (Kienesberger et al., 2009; Perry et al., 2015; Shum et al., 2016; Titchenell et al., 2016). Moreover, deficiency of hepatic fatty acid transporter CD36 reduces fat accumulation in the liver and protects against steatosis (Wilson et al., 2016). While Nrip1KO in adipocytes does not appear to affect lipolysis (Peppler et al., 2017; Tsagkaraki et al., 2023), it does enhance adipocyte fatty acid oxidation under proper stimulation and therefore might be expected to decrease levels of circulating fatty acids that can enter the liver. However, no change in serum free fatty acids is observed in the AdNrip1KO mouse (Table 1), suggesting that fatty acids may not be the conduit between Nrip1/Rip140 depleted adipose tissue and the liver.

Based on the above considerations, we hypothesized that additional effectors secreted by adipose tissues from AdNrip1KO mice mediate the observed improvements in hepatic steatosis and insulin sensitivity. To search for such effectors from adipocytes, RNAseq was performed on iWAT from Nrip1^flox/flox^ and AdNrip1KO mice and compared to RNAseq data we previously obtained from control and Nrip1KO mouse adipocytes which are implanted into mice. Unexpectedly, these RNA-seq analyses revealed that most of the upregulated genes in response to adipocyte Nrip1KO in white adipose tissue of AdNrip1KO mice encoded genes most highly expressed in macrophages and immune cells rather than adipocytes (Tables S2, S3). In particular, virtually all of the genes predicted to encode secreted proteins adipocyte-selective Nrip1KO mice were most highly expressed in adipose tissue macrophages, not adipocytes (Fig. S3 and Table S2). This observation is also supported by the markedly altered expression of the macrophage marker gene transcriptomic profile in AdNrip1KO mice (Fig. 5H-J and Table S1), accompanied by a clear shift in macrophage polarization toward the alternative M2 phenotype (anti-inflammatory) in iWAT of AdNrip1KO mice (Fig. 5H-I and Table S1).

In the context of adipose tissue browning, these results are consistent with multiple studies showing that classical thermogenic stimuli that increase Ucp1 expression also promote M2-like adipose tissue macrophage polarization. Such stimuli include chronic cold exposure (Hui et al., 2015; Yuan et al., 2024), treatment with the PPARγ agonist rosiglitazone (Rohm et al., 2024; Stienstra et al., 2008), various browning mouse models (Kumari et al., 2016; Henriques et al., 2020; Rajasekaran et al., 2019) and burn-induced browning observed in the adipose tissue of burn patients (Abdullahi et al., 2019). Thus, the results presented in this report add adipocyte Nrip1KO to the known thermogenic conditions that promote appearance of M2-like macrophages in subcutaneous adipose tissues. Based on these observations, multiple studies have demonstrated that M2 macrophages are essential for optimal cold-induced thermogenesis and energy expenditure by either enhancing sympathetic activity indirectly (Nguyen et al., 2011; Wolf et al., 2017; Wang et al., 2021) or regulating adipocyte browning directly (Henriques et al., 2020; Wu et al., 2024; Yuan et al., 2024). In this report, we complement these previous studies that showed M2-like adipose tissue macrophages signal to adipocytes by suggesting that these macrophages also serve an endocrine function by signaling to liver.

The mechanisms by which adipocytes regulate M2 macrophage polarization remain poorly understood. Various adipokines secreted by thermogenic adipocytes such as adiponectin (Hui et al., 2015), Fgf21 (Li et al., 2018) and Cxcl14 (Cereijo et al., 2018) have been reported to modulate M2 macrophage polarization. However, both *AdipoQ* and *Fgf21* gene expression were decreased in AdNrip1KO iWAT (Table S3). Also, plasma adiponectin levels were significantly reduced in AdNrip1KO mice (Table 1). *Cxcl14* expression showed no significant difference between the Nrip1^flox/flox^ and AdNrip1KO counterparts. Another potential mode of communication involves intercellular mitochondrial transfer, as several studies have proposed the possibility of mitochondria being transferred from adipocytes to neighboring macrophages, a mechanism that warrants further investigation in AdNrip1KO mice (Brestoff et al., 2021; Goodman et al., 2025; Zuo et al., 2024).

To evaluate whether adipose tissue macrophages are a main source of factors that decrease liver fat and glucose production in AdNrip1KO mice, we identified known regulators of liver steatosis among the list of genes most upregulated according to the RNAseq data. (Fig 6A). One of the top upregulated genes was the gene encoding the IL-1β antagonist IL-1rn, previously also reported to be expressed in thermogenic adipose tissue (Juge-Aubry et al., 2003; Stienstra et al., 2008). Studies have indicated that the IL-1β signaling pathway promotes hepatic steatosis through stimulation of hepatic lipogenesis (Petrasek et al., 2011, 2012; Negrin et al., 2014) and it is known that IL-1rn is a potent natural inhibitor of IL-1β (Hirsch et al., 1996; Witkin et al., 2002). We now show here that IL-1rn protein is strongly increased in the circulation of AdNrip1KO mice (Fig 6C), which likely derives from the increased ATMs in these mice. Many studies have demonstrated that IL-1rn is functional *in vivo* in alleviating hepatic dysfunction. For example, strong evidence published by our group and others that shows treatment with Anakinra—a recombinant form of IL-1rn that antagonizes IL-1 signaling via the IL-1 receptor (IL-1R)—markedly reduced obesity induced hepatic steatosis and the expression of lipogenic genes (Petrasek et al., 2012; Negrin et al., 2014). Indeed, the recombinant form of IL-1rn robustly reduces liver steatosis at blood concentrations that are similar to those reported here for IL-1rn in AdNrip1KO mice (Negrin et al, 2014). Furthermore, the specific deletion of IL-1R from hepatocytes or whole-body KO largely blocks IL-1 driven liver inflammation (Gehrke et al., 2018) and ameliorates microvascular steatosis and liver injury (Petrasek et al., 2012; Gehrke et al., 2022;). Taken together, these published results support a key role of iWAT-derived IL-1rn in modulating hepatic lipid metabolism and contributing to the beneficial systemic effects observed in AdNrip1KO mice.

Several other adipokines with metabolic regulatory functions to affect liver metabolism have been reported in brown and beige adipose tissue, including neuregulin 4 (Nrg4) (Wang et al., 2014), IL-6 (Stanford et al., 2013) and growth differentiation factor 15 (Gdf15) (Luan et al., 2019; Sjøberg et al., 2023). Notably, our RNAseq analysis of AdNrip1KO iWAT and of Nrip1KO adipose tissue implants also identified *Gdf15 among the* 61 commonly upregulated genes annotated as secreted proteins (Fig. 6A). However, circulating Gdf15 protein levels did not reach statistical differences between control and AdNrip1KO littermates under HFD conditions (Table 1). Neither was *Nrg4* mRNA expression significantly changed in AdNrip1KO iWAT and IL-6 mRNA expression was nearly undetectable in our RNAseq experiments (Table S1).

Another unexpected result reported here is the increase in masses of iWAT and gWAT in AdNrip1KO mice on HFD in spite of no change in total body weight (Fig. 2). In contrast, the masses of these adipose depots are markedly diminished in the whole body Nrip1KO mouse (Leonardsson et al., 2004). However, in both these mouse models, the sizes of adipocytes are smaller than in control mice and have an enhanced thermogenic gene expression profile (Fig. 2). These data indicate that the iWAT and gWAT adipose depots in the AdNrip1KO mice contain greater numbers of adipocytes than their Nrip1^flox/flox^ counterparts. These findings suggest the hypothesis that Nrip1 depleted adipocytes in the AdNrip1KO mice may signal to progenitors in the tissue to stimulate their differentiation into small adipocytes. The features of these adipose depots in the AdNrip1KO mouse are reminiscent of the actions of PPARγ agonists, which also cause browning and expansion of adipose tissue mass in association with their insulin sensitizing effects (Hallakou et al., 1997; Stienstra et al., 2008). Consistent with these latter reports, transgenic mice expressing PPARγ in adipocytes precursors also display adipose tissues with decreased cell size profiles (Sugii et al., 2009; Shao et al., 2018, 2021). Nrip1/Rip140 is known to interact with and suppress PPARγ transcriptional activity in addition to many other nuclear receptors, suggesting a mechanism to explain the effects of Nrip1 depletion to decrease adipocyte size in our AdNrip1KO mice. Similar observations have been made when PPAR is activated in mice and humans (Tontonoz et al., 1998; Meier et al., 2002; Stienstra et al., 2008;). Interestingly, a recent study has showed that transient pharmacological inhibition of IL-1 signaling with sIL-1rn prompts adipocyte proliferation in both iWAT and gWAT (Hofwimmer et al., 2024). It will be important to direct future studies towards identifying whether this or other immune cell factors modulate progenitor cell differentiation in response to Nrip1KO.

Also noteworthy in our study is the decrease in serum cholesterol and LDL levels in AdNrip1KO mice in spite of normal serum free fatty acid levels and total triglycerides (Table 1). This finding is of particular significance in light of the associations between polymorphisms in the NRIP1 gene locus and cardiovascular, hepatic and blood triglyceride phenotypes (https://hugeamp.org/gene.html?gene=NRIP1) (Dornbos et al., 2022). The decreases in serum total cholesterol and LDL levels in particular are of potential significance in cardiovascular disease, highlighting the interesting question of whether these effects of adipose Nrip1 depletion may extrapolate to beneficial effects in human atherosclerosis.

The key difference between the whole body Nrip1 null mouse (Leonardsson et al., 2004; Seth et al., 2007) and the adipose selective AdNrip1KO mouse model revealed in this study is the lack of differences in body weight (Fig. 2C) and oxygen consumption rates (Fig. 4A) upon comparing the AdNrip1KO mice to wild type mice. Neither did AdNrip1KO mice show any divergence from Nrip1^flox/flox^ mice in RER values throughout the day or night (Fig. 4B). One caveat in these studies is that they were carried out at room temperature, and we did detect a small difference in the ability of AdNrip1KO mice to better maintain body temperature after several hours of cold exposure when fasting (Fig. 1I). This indicates some increase in energy expenditure in AdNrip1KO mice in the form of heat production under cold stress conditions, but not at elevated temperatures. Nonetheless, the dramatic enhancement of energy expenditure in Nrip1 null mice at room temperature (Seth et al., 2007) is not replicated in the AdNrip1KO mice (Fig. 4). This indicates one or more other tissues that express Nrip1/Rip140 contribute to the energy expenditure phenotype of Nrip1 null mice. Skeletal muscle is an obvious candidate for such a contribution since this tissue derived from either the whole body Nrip1KO mouse (Leonardsson et al., 2004) or the striated muscle-selective Nrip1KO mouse exhibits strong upregulation of expression of genes associated with catabolic activity (Yamamoto et al., 2023), including those encoding enzymes of fatty acid oxidation pathways. This hypothesis that muscle metabolism under Nrip1KO conditions contributes largely to increased systemic energy expenditure also would explain why the adipose tissue depots in Nrip1 null mice contain less fat compared to the AdNrip1KO mice under conditions of similar food intake. It is also possible that other tissues that express Nrip1/RIP140 contribute to the increased systemic energy expenditure of whole body Nrip1KO mice.

In summary, the present study demonstrates that specific deletion of Nrip1 in adipose tissue can account for the alleviation of glucose intolerance, hepatic steatosis and insulin resistance previously observed in whole body Nrip1KO mice. However, the AdNrip1KO model does not replicate the large increase in energy expenditure observed in Nrip1 null mice, suggesting other tissues are involved in this response. Importantly, our findings suggest a novel adipocyte–M2 macrophage derived IL-1rn–liver axis that contributes to systemic metabolic regulation, highlighting a key hypothetical endocrine role of ATMs. A similar involvement of M2-like adipose tissue macrophages has been suggested related to small extracellular vesicles that may contribute to rosiglitazone-induced insulin sensitization (Rohm et al., 2024). From a therapeutic standpoint, targeting adipose tissue Nrip1/Rip140 has the potential to be an effective approach to treat insulin resistance, type 2 diabetes and fatty liver disease.

## Methods

### Animal Husbandry

All animal studies performed were approved by the Institutional Animal Care and Use Committee (IACUC) of the University of Massachusetts Chan Medical School. Unless otherwise indicated, all mice were housed in ventilated cages (3-5 mice/cage) with an automated watering system, 12 hour-Light/12 hour-Dark cycle, enrichment, and normal mouse chow diet (LabDiet, 5P76 Prolab® RMH 3000) were given *ad libitum*.

### Mice Generation and Genotyping

Nrip1^flox/flox^ mice were developed at Cyagen Bioseciences (US) by introducing two specific sgRNA sequences that spanned exon 4 of *Nrip1* gene (Fig.S1A). The gRNAs to *Nrip1* gene, the donor vector containing loxP sites, and Cas9 mRNA were co-injected into fertilized mouse eggs to generate targeted edited offspring. F0 founder animals were identified by PCR and Sequencing. F0 founder mice were bred to wild-type mice to test germline transmission and produced F1 offspring. Adipose specific Nrip1 knockout (AdNrip1KO) mice were generated by crossing the Nrip1^flox/flox^ mice to Adiponectin-Cre mice, B6; FVB-Tg (Adipoq-cre)1Evdr/J (Jackson Labs, 010803). Nrip1^flox/flox^ were used as control animals to compare to AdNrip1KO mice in our studies.

Genomic DNA was isolated from ear punches at 21-days old or younger pups. Ear samples were digested with 100 μL lysis buffer (50 mM Tris pH 8.0, 50 mM kCl, 2.5 mM EDTA, 0.45% NP40, 0.45% Tween 20, 100 μg/mL proteinase K) at 65 ^°^C overnight, followed by 95 ^°^C inactivation for 10 min. Genotyping was performed by PCR analysis. For *Nrip1* gene genotyping, the following specific oligonucleotide primers were used, forward primer: 5’-ATGTCATTCCACAGTGCTGAAAC -3’, reverse primer: 5’-TTTCCTAAATCAGAACCCACCCC-3’. For Adiponectin-Cre mice genotyping, the primers used were forward primer: 5’-CTAGGCCACAGA ATTGAA AGATCT-3’ and reverse primer: 5’-CTAGGCCACAGAATTGAAAGATCT-3’. All PCR reactions were carried out in 20 μL volume system using KAPA Mouse Genotyping Kits (Roche, 07961316001), 10 μM primers and 1 μL DNA template.

### Acute cold exposure

Twenty-week-old male mice on a chow diet were singly housed and placed in an environmental chamber at 6 °C for 6 hours without food, but with free access to water. Body weight and body temperature was monitored hourly, the latter of which was measured using a rectal probe (Braintree Scientific, RET-3). At the end of the experiment, all mice were brought back to room temperature with access to food and water *ad libitum*.

### High-Fat-Diet feeding (HFD)

For HFD feeding experiment, twelve-week-old male control Nrip1^flox/flox^ mice and AdNrip1KO mice were fed *ad libitum* with irradiated HFD (D12492i, 60% from fat, Research Diet Inc.) for up to 16-weeks at room temperature.

### Glucose Tolerance Test

Mice were fasted overnight for 16-hours (5.00pm-9.00am), with free access to drinking water. Next morning, body weights were measured, and blood samples was taken from the tails to quantify their basal blood glucose using a Contour Next Blood Glucose Meter. Afterwards, D (+) glucose (Sigma-Aldrich) was administered at 1 g/kg body weight by intraperitoneal injection. Sequentially, blood glucose levels were measured from the tail vein at 15, 30, 60, 90 and 120 min post glucose injection. After the glucose tolerance test, all mice were placed back in clean cages with free access to food and water.

### Insulin Tolerance Test

Mice were fasted for 4-hours with free access to water. Body weights were measured, and blood samples were taken from the tails to quantify their basal blood glucose using a Contour Next Blood Glucose Meter. After injection of 1.5 U/kg human insulin, blood glucose levels were measured at 15, 45, 60, 90 and 120-min using a Contour Next Blood Glucose Meter. Immediately following glucose tolerance testing, mice were placed back in clean cages with free access to food and water.

### Hyperinsulinemia-euglycemic clamp to assess insulin sensitivity

The hyperinsulinemic-euglycemic clamp experiments were performed at the Metabolic Disease Research Center of the University of Massachusetts Chan Medical School as previously described(Friedline et al., 2024). The insulin clamps were conducted in Nrip1^flox/flox^ and AdNrip1KO mice after 16 weeks of HFD. Briefly, an indwelling jugular catheter was placed during survival surgery. Approximately 5 days later, mice were fasted for 14 hours, followed by a continuous basal glucose infusion of D-[3-^3^H]glucose at 0.05 uCi/min. At the end of the basal period 20 μL of blood was taken for whole-body plasma glucose turnover and hepatic glucose production and will be used as a basal measurement (Friedline et al., 2024). Following the basal glucose infusion, insulin was delivered in a single bolus at 150 mU/kg, followed by a continuous infusion of 2.5 mU/kg/min (Novolin R; Novo Nordisk, Denmark) to raise plasma insulin to a physiological level of 0.8 ng/mL (Friedline et al., 2024). Then, 10μL of blood was collected at 10-20 min intervals for plasma glucose measurements and a 20% dextrose was infused at variable rates to maintain euglycemia at 120 mg/dL. During the clamp, 0.1 μCi/min [3-^3^H]glucose was infused continuously to measure whole body glucose turnover, glycolysis, glycogen plus lipid synthesis and hepatic glucose production during the insulin stimulated state (Friedline et al., 2024). To measure hepatic insulin action, the percent change from basal HGP to insulin-stimulated clamp HGP was compared. Insulin-stimulated glucose uptake in organs was measured by infusing a bolus of 10 μCi 2-[1-^14^C]deoxy-D-glucose (2-[^14^C]DG) at 75 min post clamp start. Blood samples were taken at 80, 85, 90, 100, 110 and 120 min after clamp start to measure [^3^H]glucose, ^3^H_2_O, and 2-[^14^C]DG concentrations. At the end of the clamp study, tissues were taken for biochemical and molecular analysis such as liver, adipose, and skeletal muscle from the mice while under anesthesia (Friedline et al., 2024).

### Energy balance analysis using metabolic cages

After 14 weeks of HFD, indirect calorimetry and energy balance parameters, including food/water intake, energy expenditure, respiratory exchange ratio, and physical activity, were noninvasively measured for 3 days using metabolic cages with an environmental chamber (TSE-Systems Inc., Chesterfield, MO) (Friedline et al., 2024). Respiratory exchange ratio (RER) and energy expenditure rates were calculated using an online software CalR (https://bankslab.shinyapps.io/prod_ver/), while the total body mass was used as a selective covariant (Mina et al., 2018).

### RNA isolation and RT-qPCR

Total RNA was isolated from mouse tissues by homogenizing tissue in TRIzol Reagent (Invitrogen, 15596018) with a stainless-steel bead in a microfuge tube using a Tissue Lyzer II (Qiagen, 85300), following the manufacturer’s instructions for RNA purification. cDNA was synthesized using iScript cDNA Synthesis Kit (BioRad, 1708891). QPCR were performed on a BioRad CFX96 thermocycler using an iTaq Universal Sybr Green Supermix (BioRad, 1725120). Primer sequences are provided in Table 2. The relative mRNA expression was analyzed with the ΔΔCT method.

**Table 2.**
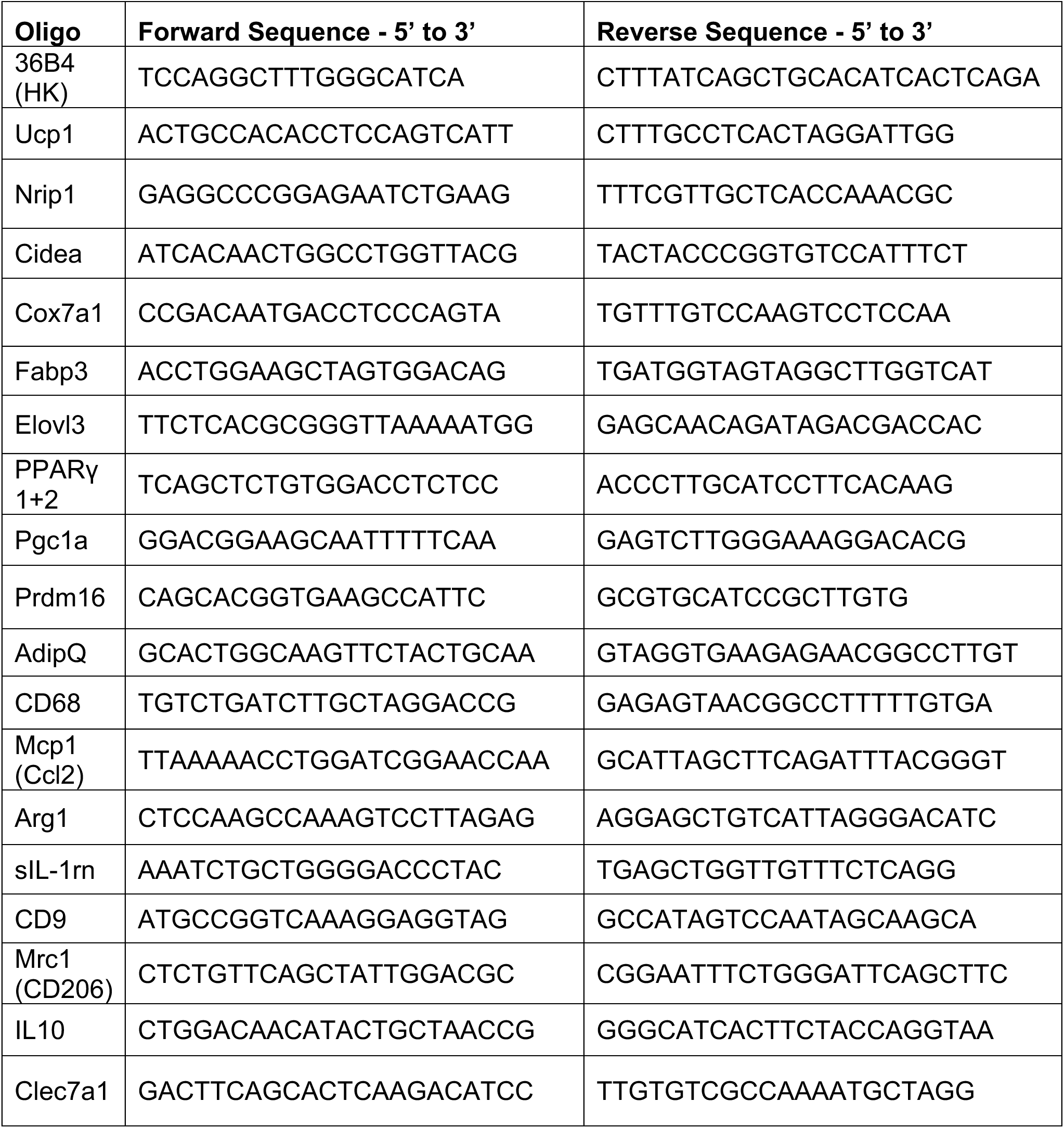
Primers used for RT-qPCR.

### RNA-sequencing

Total RNA was isolated from mouse tissues as above, 2 μg of isolated RNA were submitted to GENEWIZ for total RNA sequencing. The data were acquired demultiplexed in the form of FASTQ files. Alignment and quantification of gene expression levels were performed using the DolphinNext RNA-seq pipeline(Yukselen et al., 2020). DEBrowser was then used on these expression levels to find differentially expressed genes between two genotypes, the padj was set to 0.05 and a minimal fold change (minFC) =1.5 or 2(Kucukural et al., 2019). Principal component analysis was applied to the normalized data (DESeq2 getNormalizedMatrix with method = “MRN”). Once DE genes were found, enriched pathways were identified using the enrichGO-biological process (GO-BP) and enrichKEGG pathway routine with *p* value Cutoff of 0.05. Heatmaps of selected genes were generated using the normalized (for sample depth) values returned by the DESeq2 counts (ddsRES, normalized = TRUE) function. The differentially expressed genes list were shown in Supplementary Table 1 (Table S1).

### Western Blots

For tissue protein expression analyses, mouse tissues were prepared following protocol as previously described (An & Scherer, 2020). Protein concentrations from sample tissue were quantified with the Bicinchoninic Acid (BCA) assay (Pierce, A65453) following manufacturer’s directions. Protein samples were run on Protein TGX Precast Protein Gels (BioRad, 5678085), followed by transferring to nitrocellulose using Trans-Blot Turbo Mini Nitrocellulose Transfer Pack (BioRad, 1704158) and the Trans-Blot Turbo Mini System (BioRad, 1704159). Protein transfer verification was performed by incubating membranes with Ponceau solution (0.1% wt/v Ponceau S, 5% v/v Glacial acetic acid in water) for 3 min with gentle shaking, followed by rinsing with water to remove excess ponceau stain. Membranes were blotted with the primary antibodies in Table 3.

**Table 3.** Reagents and resources.

| **REAGENT for Immunoblotting** | **SOURCE** | **IDENTIFIER** |
| --- | --- | --- |
| Rabbit monoclonal Ab-Ucp1<br>1:500 dilution | Abcam | Cat # ab209483 |
| Rabbit Phospho-Akt (Ser473) Antibody<br>1:1000 dilution | Cell Signaling | Cat # 9271 |
| Rabbit total Akt Antibody<br>1:1000 dilution | Cell Signaling | Cat # 9272 |
| Rabbit mAb-Hsp90 (C45G5) (HRP<br>Conjugate)<br>1:1000 dilution | Cell Signaling | Cat # 79641 |
| Total OXPHOS Rodent WB Antibody<br>Cocktail<br>1:1000 dilution | Abcam | Cat # ab110413 |
| Mouse monoclonal Ab-Tubulin<br>1:1000 dilution | Sigma-Aldrich | Cat # sc5286 |
| Goat Anti-Rabbit IgG-HRP<br>1:10000 dilution | Invitrogen | Cat # 65-6120 |
| Goat Anti-Rabbit IgG-HRP<br>1:10000 dilution | Invitrogen | Cat # 65-6520 |
| **REAGENT for Immunohistochemistry** | **SOURCE** | **IDENTIFIER** |
| Rabbit monoclonal Ab-Ucp1<br>1:50 | Abcam | Cat # ab209483 |
| Rabbit Anti-CD68 antibody<br>1:50 dilution | Abcam | Cat # ab125212 |
| Goat anti-Rabbit-IgG-Alexa594<br>1:500 dilution | Thermofisher | A11012 |
| **RESOURCES** | **SOURCE** | **IDENTIFIER** |
| Mouse Insulin ELISA Kit | Invitrogen | Cat # EMINS |
| Free Fatty Acid Fluorometric Assay Kit | Cayman | Cat # 700310 |
| mouse IL-1Ra/IL-1F3 ELISA Kit | Biotechne,<br>R&D system | Cat # MRA00 |
| Mouse & Rat GDF-15 ELISA Kit | Biotechne,<br>R&D system | Cat # MGD150 |
| Mouse Adiponectin ELISA Kit | RayBiotech | Cat # ELM-Adiponectin<br>ACRP30 |
| Triglyceride determination kit | Sigma-Aldrich | Cat # TR0100 |

Next morning, membranes were washed with 0.1% TBS-T for 3 times, 5 min per wash. Secondary HRP Antibodies were diluted 1:5000 in 0.1% TBS-T with 5% wt/v BSA and incubated for 1-hour at room temperature, with gentle shaking. Membranes were washed with 0.1% TBS-T, followed by incubating with enhanced chemiluminescence (ECL) (Perkin Elmer, NEL104001EA) for detection using a Bio-Rad Chemi-Doc.

### Histological and Immunohistochemical analysis of tissues samples

For the immunohistochemistry, tissue samples were fixed in 4% paraformaldehyde and embedded in paraffin. Sectioned slides were then stained at UMass Chan Medical School Morphology Core for Hematoxylin and Eosin (H&E) staining and for immunohistochemistry. For immunohistochemistry, Anti-Ucp1 antibody (Abcam, ab209483) or Anti-CD68 antibody (Abcam, ab125212) was incubated at 1:50 (Table 3). Immunohistochemistry slides were analyzed either with an Axiovert 35 Zeiss microscope (Zeiss) equipped with an Axiocam CCl camera or with a Leica DM2500 LED equipped with a Leica MC170 HD using a 5x/0.12 objective or a 20x/0.4 Leica objective. Images were manually adjusted for threshold in Fiji (ImageJ). The eraser function was then used to manually further separate cells which remained incorrectly attached to each other. “Analyze particles” with a minimum size of 10 pixels and excluding cells which crossed the image boundary was then used to count the cells and measure their area (in pixels).

### Immunofluorescent staining of adipocytes *in vitro*

For the immunofluorescence, adipocytes on coverslips were washed 3 times with 1xPBS, then fixed with 4% paraformaldehyde in PBS for 10-15 min at room temperature, followed by washing in 1x PBS. Cells on coverslips were blocked with Normal goat serum buffer (NGSB) (2% v/v goat serum, 1%w/v BSA, 0.1% v/v Triton X-100, 0.05% v/v Tween-20 in 1x PBS) for 1-hour at room temperature. Primary antibody Rabbit Anti-Ucp1 (Abcam, ab209483) diluted at 1:100 in NGSB buffer, was incubated overnight at 4 °C. After primary antibody incubation, the cells were washed three times with 1x PBS, followed by incubating with fluorescently labeled secondary antibody, Goat anti-Rabbit-IgG-Alexa594 1:500 (Thermofisher, A11012) for 1-hour at room temperature. Coverslips were washed three times with 1x PBS. LipidTOX^TM^ Green (Invitrogen, H34475) was diluted in PBS at 1:200 and incubated with the coverslips for 30 min at room temperature, followed by washing three times with 1x PBS. Nuclei were stained for 5 min with 1 ug/mL DAPI in PBS. Coverslips were mounted in Prolong Glass Antifade reagent (Invitrogen, P36982).

### Liver triglyceride content quantification

Liver samples (50-200 mg) were first saponified in ethanolic KOH, following a protocol described previously (Jouihan, 2012). After the tissue lysates were neutralized with MgCl_2_, tissue lysates were kept on ice for 10 min, and centrifuged at 10,000 x g for 5 min. The supernatant was removed for glycerol content quantification, using Sigma Triglyceride determination kit (Sigma-Aldrich TR0100) following the manufacturer’s instruction.

### Plasma insulin, free fatty acids, total triglyceride, total cholesterol, HDL and LDL IL-1rn, Gdf15, Adiponectin, Leptin quantifications

Plasma insulin content was quantified using Mouse INSULIN ELISA Kit (Invitrogen, EMINS) according to the manufacturer’s instruction. Plasma Free Fatty Acid was measured using Free Fatty Acid Fluorometric Assay Kit (Cayman, No. 700310) according to the manufacturer’s instruction. Plasma triglyceride, total cholesterol, HDL were analyzed using Cobas Analyzer (Roche Diagnostics, Indianapolis, IN, USA) at University of Massachusetts Chan Medical School Metabolic Disease Research Center as described in previously studies (Caracciolo et al., 2018). Plasma LDL content was quantified as: LDL = TC/0.948 – HDL/0.971 – (TG/8.56 + [TG x NonHDL]/2140 – TG2/16100) – 9.44 (Sampson et al., 2020). Plasma VLDL was quantified as: VLDL=TG/5. The plasma IL-1rn (Biotechne, R&D system, mouse iL-1Ra/iL-1F3), Gdf15 (Biotechne, R&D system, MGD150), Adiponectin (RayBiotech, ELM-Adiponectin), Leptin (RayBiotech, ELM-Leptin) were quantified according to the manufacturer’s instruction.

### *Ex vivo* fat tissue glucose uptake assay

Inguinal, gonadal, and brown adipose tissue fat pads were harvested from Nrip1^flox/flox^ and AdNrip1KO male mice after 16 weeks high-fat diet feeding. The fat pads were weighed and then minced into small pieces and equally divided into 2 wells of a 24-well tissue culture plate with 1 mL of glucose-free DMEM (Invitrogen, A1443001). Once all tissue was placed in wells, 50 μL of 100 mM D (+)-glucose (Sigma, G5146) was added to each well for a final concentration of 5 mM glucose. Plates were covered and placed in a 37°C cell incubator with 5% CO_2_ and 10 uL of culture media was removed at 2, 4, 6 and 8-hours from each well and placed in tubes at -20°C overnight. To quantify the glucose, tubes with 10 uL of media were thawed and allowed to come to room temperature the next day. For glucose measurements, a GlucCell Cell Culture Glucose Monitoring System (ChemGlass Life Sciences, CLS132202) was used to quantify the glucose concentration left in the culture media of each time point. Tubes containing media were gently mixed and 1.5 μL of media was placed on the GlucCell strip to determine the amount of glucose in media at each timepoint. Wells containing only 5 mM glucose in media without tissue were used as baseline controls at the starting point and labeled as Glucose concentration at 0 hour. Glucose uptake was quantified as follows: Glucose uptake= ((Glucose Concentration at 0 hour) - (Glucose Concentration at 2,4,6,8-hour)) / tissue mass.

### Primary white adipocyte preparation and differentiation

Inguinal WAT was dissected from 5-week-old male and female mice. The primary adipocytes were isolated and cultured as described previously (Henriques et al., 2020; Tsagkaraki et al., 2021, 2023).

### Statistical analysis

All statistical analysis for the data was performed in GraphPad Prism version 9.0. Statistical tests performed are listed in each figure legend. Data are presented as means ± SEM. ns, not significant, \**p* < 0.05, \*\**p* < 0.005, \*\*\**p* < 0.0005, \*\*\*\**p* < 0.0001.

## Authors Contributions

H.W. and M.P.C. designed the study, interpreted the data and wrote the manuscript. H.W performed most of the experiments and analyzed the data. M.K. established the mouse colonies, performed the genotyping and the collected blood samples from the mice. N. A. performed the qPCR validation in Figure 5I. E.T. and S.M.N performed the implantation study and prepared the samples for Nrip1KO-implantation RNA sequencing. S.M.N helped to write the methods sections of the manuscript. L.M.L. supported the quantification of adipocyte size and the RNA sequencing data analysis. A.G offered technical support for quantifying *ex vivo* glucose uptake in adipose tissues. N. A., S.M.N, L.R., K.B.S., B.Y., N.R., all helped to collect tissue samples from mice. SC and TD collaborated in studies on adipose tissue implantations. M.S., H.C., and L.A.T. performed the metabolic cage and hyperinsulinemic-euglycemic clamp studies and analyzed the metabolic data. J.K.K. designed the metabolic cage and hyperinsulinemic-euglycemic clamp studies, analyzed and interpreted the data, and edited the manuscript.

## Acknowledgements

We thank Czech laboratory members for their excellent discussions related to this project. We thank the Morphology Core at UMass Chan Medical School for performing the histological and immunochemical staining of the tissue samples. We thank Dr. William Theuerkauf and Birgit Koppetsch at UMass Chan Medical School, Program in Molecular Medicine for assistance with confocal microscopy. We thank Dr. Roger Davis and Dr. Norman Kennedy at UMass Chan Medical School, Program in Molecular Medicine for assistance with the Axiovert 35 Zeiss microscope. This work was supported by National Institutes of Health grant DK130852 (to M.P.C.) and DK133772 (to J.K.K.). We also gratefully acknowledge generous funding through the Isadore and Fannie Foxman Chair in Medical Science (to M.P.C.).

**The authors declare no conflicts.**

## Figure Legends

**Supplementary Figure 1:**
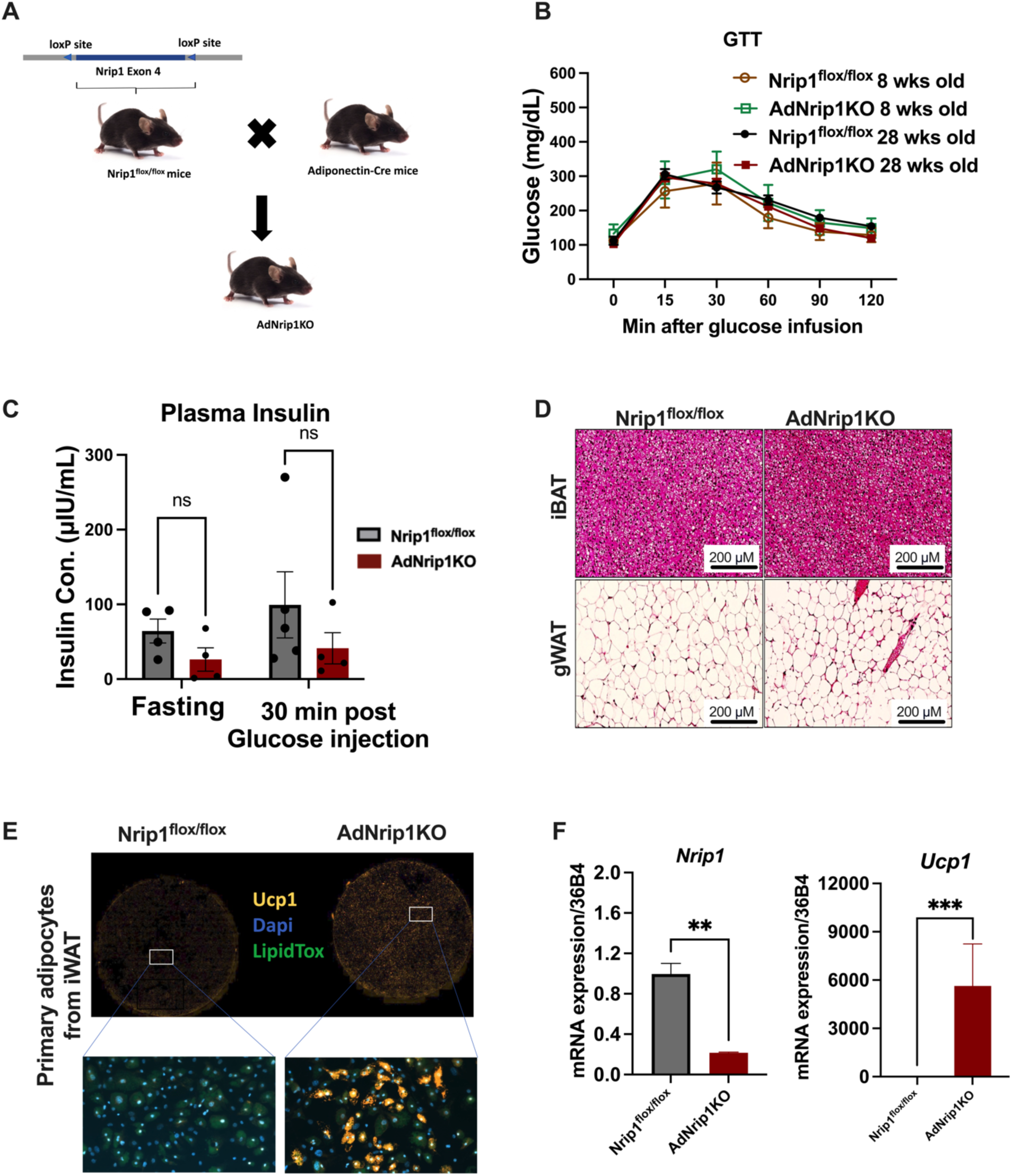
Generation of AdNrip1KO mice. **A.** Schematic illustration of the generation of AdNrip1KO mice: Nrip1^flox/flox^ mice were generated by introducing two specific sgRNA sequences spanning exon 4 of *Nrip1* gene. The AdNrip1KO mice were generated by crossing the generated Nrip1^flox/flox^ mice to Adiponectin-Cre mice. **B.** Blood glucose from GTT of 8-week-old and 28-week-old mice fed on chow diet (n = 5-7 mice for each genotype). **C.** Plasma insulin concentration from 12-week-old Nrip1^flox/flox^ and AdNrip1KO mice fed on chow diet. **D.** Representative Hematoxylin & Eosin staining of iBAT and gWAT isolated from the 12-week-old Nrip1^flox/flox^ and AdNrip1KO. **E.** Immunohistochemical staining of primary adipocyte cell cultures in vitro isolated from 5-week-old mice. The slides were stained with Ucp1 antibody, LipidTox was used for detecting lipid droplets and 4′,6-diamidino-2-phenylindole (DAPI) for Nuclei. **F**. *Nrip1* and *Ucp1* relative mRNA expression from the primary adipocytes. Data are represented as mean ± SEM. In panel **B,** significance analysis was compared using Two-way ANOVA with Sidak’s multiple comparisons test. **C** was compared with One-way ANOVA with Tukey’s multiple comparisons test. **F** was compared through unpaired 2-tailed *t*-test between Nrip1^flox/flox^ vs AdNrip1KO animals. \*\**p* < 0.005, \*\*\**p* < 0.0005.

**Supplementary Figure 2:**
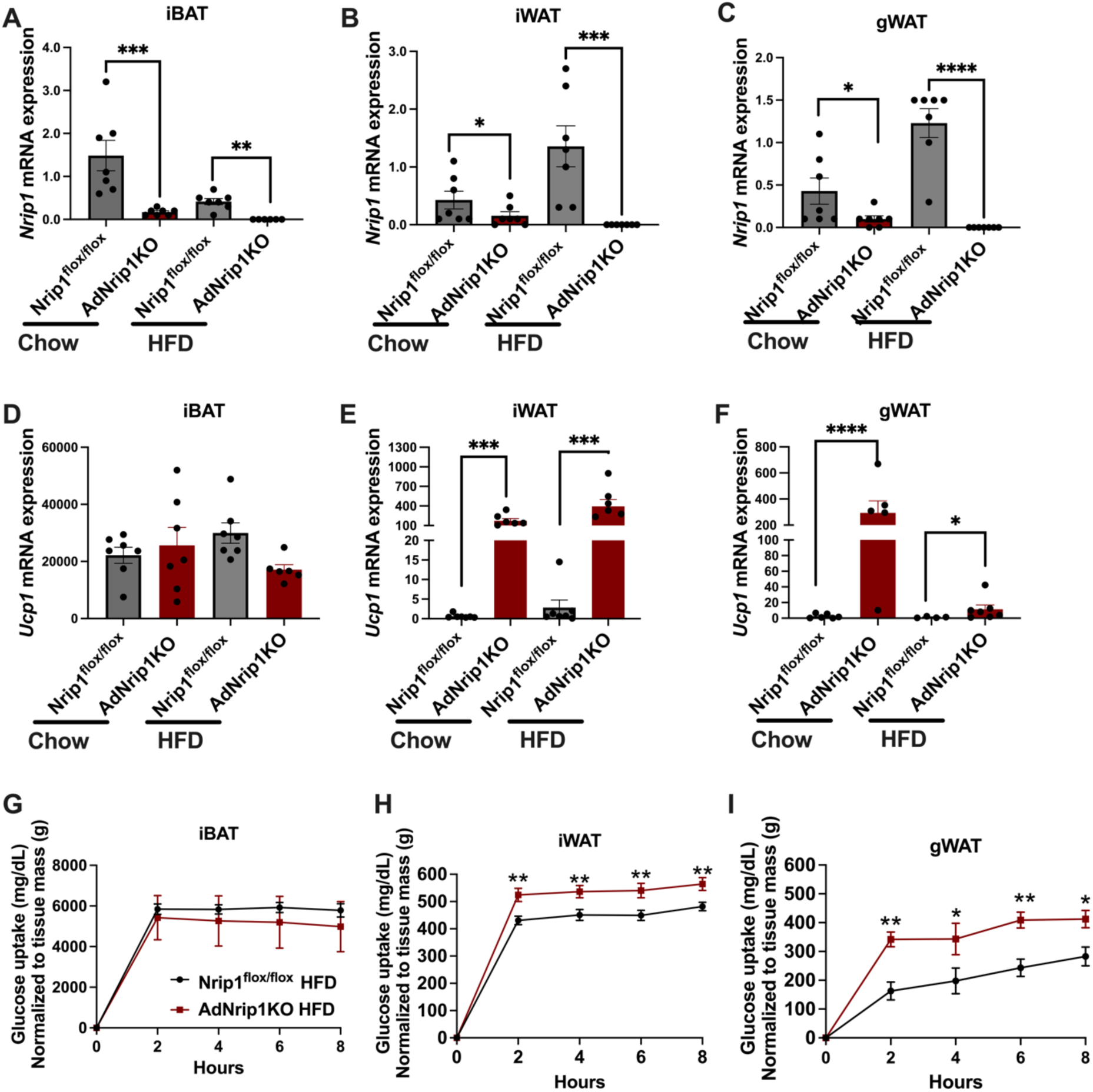
Validation of Nrip1 deficiency in Nrip1 KO mice under conditions of HFD feeding. **A-C.** *Nrip1* mRNA expression in iBAT, iWAT and gWAT of male mice on chow diet and after 16 weeks HFD feeding. **D-F.** *Ucp1* mRNA expression in iBAT, iWAT and gWAT of mice on chow diet and after 16 weeks HFD feeding. **G-I.** *Ex vivo* quantification of glucose uptake from iBAT, iWAT and gWAT of mice after 16 weeks HFD feeding. Net glucose uptake was normalized to the analyzed tissue mass. Data are represented as mean ± SEM. From **A-F**, statistical analyses were compared with by One-way ANOVA with Tukey’s multiple comparisons test. **G-I**, significance analysis was performed using Two-way ANOVA with Sidak’s multiple comparisons test. \**p* < 0.05, \*\**p* < 0.005, \*\*\**p* < 0.0005, \*\*\*\**p* < 0.0001.

**Supplementary Figure 3:**
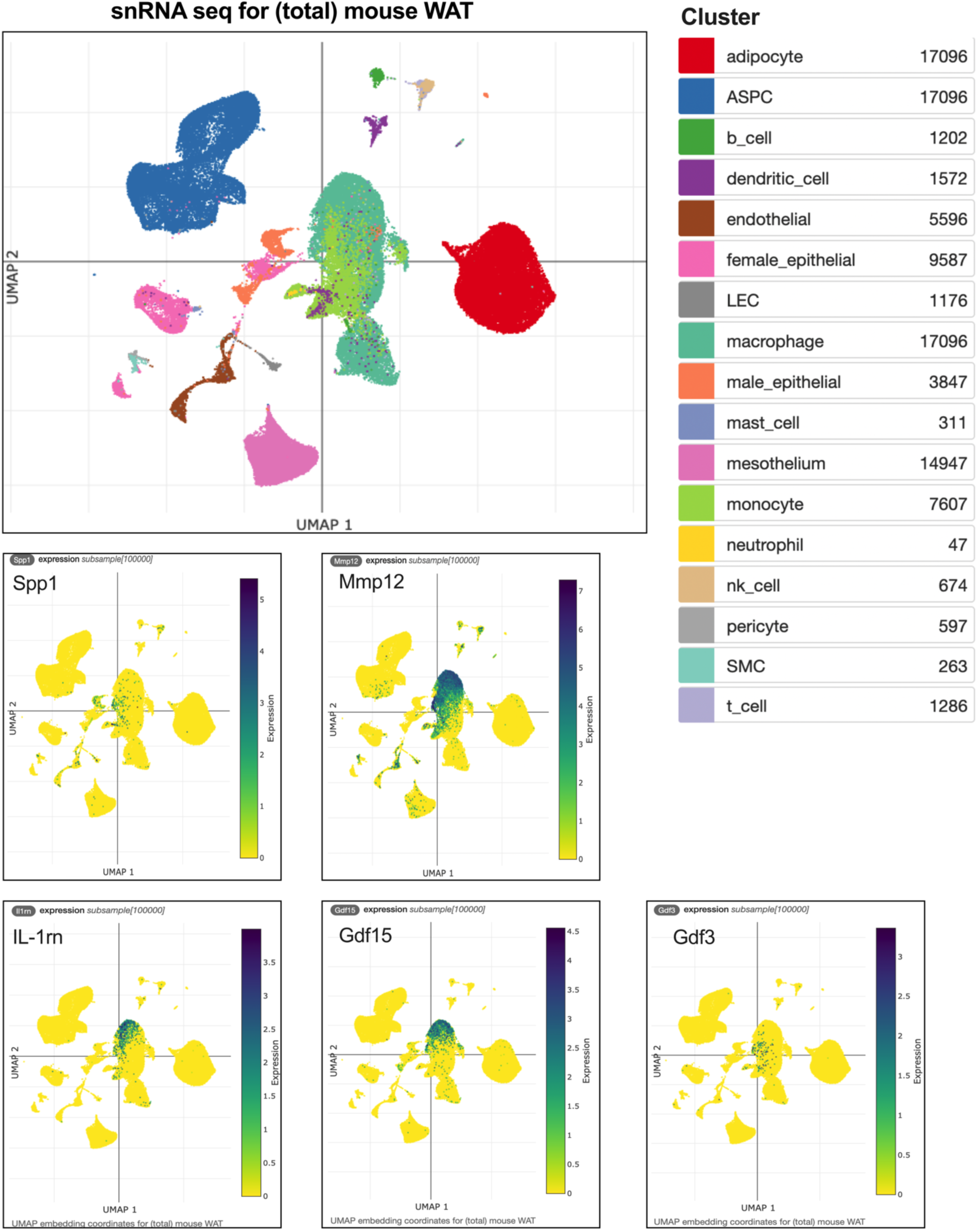
snRNAseq analysis of gene expression pattern in mouse WAT. Genes expression pattern in various cell types within total mouse WAT from the single-cell portal (Emont et al., 2022) : https://singlecell.broadinstitute.org/single_cell/study/SCP1376/a-single-cell-atlas-of-human-and-mouse-white-adipose-tissue?genes=Il1rn&cluster=Mouse%20WAT&spatialGroups=--&annotation=cluster--group--study&subsample=100000&tab=distribution The top 5 most highly upregulated genes annotated for encoding secreted proteins identified from both implanted Nrip1KO and AdNrip1KO mice were analyzed from the single cell portal.

**Supplementary Table 1: Upregulated differentially expressed genes in iWAT of AdNrip1KO mice.**

**Supplementary Table 2: Common upregulated genes predicted to encode secreted proteins from implanted Nrip1KO and AdNrip1KO mice.**

**Supplementary Table 3: Common downregulated genes predicted to encode secreted proteins from implanted Nrip1KO and AdNrip1KO mice.**

